# Detection of aberrant splicing events in RNA-seq data with FRASER

**DOI:** 10.1101/2019.12.18.866830

**Authors:** Christian Mertes, Ines Scheller, Vicente A. Yépez, Muhammed H. Çelik, Yingjiqiong Liang, Laura S. Kremer, Mirjana Gusic, Holger Prokisch, Julien Gagneur

## Abstract

Aberrant splicing is a major cause of rare diseases, yet its prediction from genome sequence remains in most cases inconclusive. Recently, RNA sequencing has proven to be an effective complementary avenue to detect aberrant splicing. Here, we developed FRASER, an algorithm to detect aberrant splicing from RNA sequencing data. Unlike existing methods, FRASER captures not only alternative splicing but also intron retention events. This typically doubles the number of detected aberrant events and identified a pathogenic intron retention in *MCOLN1*. FRASER automatically controls for latent confounders, which are widespread and substantially affect sensitivity. Moreover, FRASER is based on a count distribution and multiple testing correction, reducing the number of calls by two orders of magnitude over commonly applied z score cutoffs, with a minor sensitivity loss. The application to rare disease diagnostics is demonstrated by reprioritizing a pathogenic aberrant exon truncation in *TAZ* from a published dataset. FRASER is easy to use and freely available.

## Main

It is estimated that between 15 and 60% of the variants causing rare diseases affect splicing.^1–3^ The mechanisms include skipping, truncation, and elongation of exons as well as intron retention.^4,5^ Despite advances in detecting variants affecting splicing by machine learning,^6–9^ accurate detections remain limited in particular for deep intronic variants.^9^ Therefore, genetic diagnosis guidelines require additional functional evidence to classify a variant as pathogenic.^10,11^ Further, many variants affecting splicing, especially deep intronic variants, are ignored by most prediction tools,^12^ or are missed when whole exome sequencing or panel sequencing technologies are used.^13^ To overcome the limitation of genetic variant interpretation, RNA sequencing (RNA-seq) has gained popularity over the last years.^14–17^ RNA-seq does not only allow validating or invalidating effects on splicing of variants of unknown significance,^14^ but also allows detecting *de novo* aberrant splicing events transcriptome-wide, including the activation of deep intronic cryptic splice sites.^14,15,17^

Three distinct methods developed by (i) Cummings et al.,^14^ (ii) Kremer et al.,^15^ and (iii) Frésard et al.^16^ have been employed to call aberrant splicing in RNA-seq data for rare disease diagnostics. All methods make use of the so-called RNA-seq split reads whose ends align to two separated genomic locations of the same chromosome strand and are therefore evidence of splicing events. These methods all consider RNA-seq split reads *de novo*, i.e. beyond annotated splice sites, because the creation of novel splice sites has a strong pathogenic potential by leading to frameshift, ablation of protein sequences, or creation of non-functional protein sequences. The first method consists of a combination of cutoffs applied to absolute and relative RNA-seq split read counts.^14,17^ The limitation of this method is that statistical significance is not assessed. Furthermore, the cutoffs are not data-driven. In particular, it is unclear whether requiring that an intron occurs in no other sample^14^ or in less than 5 affected samples^17^ would generalize well to larger cohorts than the ones investigated so far. Kremer et al.^15^ instead tested the significance of differential splicing using Leafcutter,^18^ a multivariate count fraction model developed for mapping splicing quantitative trait loci. This approach allowed controlling for false discovery rate and was less dependent on cohort size. One limitation, however, was made evident by Frésard et al.,^16^ who showed that strong covariations of split read based splicing metrics are widespread within RNA-seq compendia. The origins of these covariations may include sex, population structure, or technical biases such as batch effects or variable degree of RNA integrity. Not controlling for these latent confounders can substantially affect the sensitivity of detecting aberrant splicing events. To address this issue, Frésard et al. corrected split read based splicing metrics by regressing out principal components. Aberrant splicing events were then identified using a cutoff on z scores (|*z*| ≥ 2) of these corrected splicing metrics. The drawback of this approach is that an absolute z score cutoff does not guarantee any control for false discovery rate. Moreover, a z score cutoff amounts to a quantile cutoff when assuming that the data distribution is approximately Gaussian. However, Gaussian approximations may be inaccurate when splicing metrics are based on low split read counts, which occurs on splice sites with low coverages and at repressed splice sites.

Here, we address these issues with FRASER (Find RAre Splicing Events in RNA-seq), an algorithm that provides a count-based statistical test for aberrant splicing detection in RNA-seq samples, while automatically controlling for latent confounders (Figure 1). Unlike previous methods, FRASER is not limited to alternative splicing but also captures intron retention events by considering non-split reads overlapping donor and acceptor splice sites. The parameters are optimized for recalling simulated outliers by training a so-called denoising autoencoder.^19^ FRASER shows substantial improvements against former methods on simulations based on the healthy cohort dataset of the Genotype-Tissue Expression project (GTEx).^20^ Lastly, we demonstrate the usage of FRASER for rare disease diagnosis by reanalyzing an RNA-seq dataset of individuals affected by a rare mitochondrial disorder.^15^

**Figure 1.**
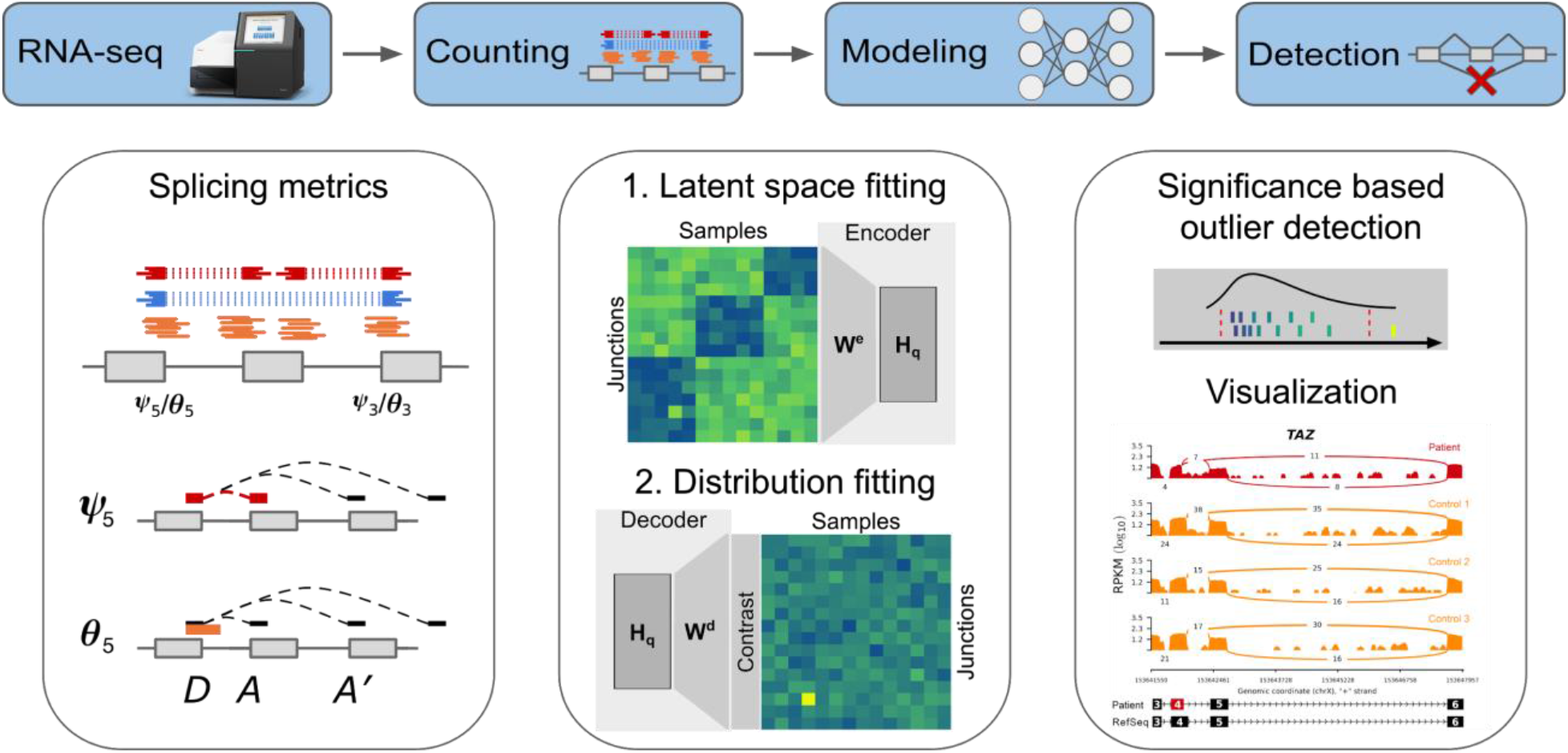
The FRASER aberrant splicing detection workflow. The workflow starts with RNA-seq aligned reads and performs splicing outlier detection in three steps. First (left column), a splice site map is generated in an annotation-free fashion based on RNA-seq split reads. Split reads supporting exon-exon junctions as well as non-split reads overlapping splice sites are counted. Splicing metrics quantifying alternative acceptors (*ψ*_5_), alternative donors (*ψ*_3_) and splicing efficiencies at donors (*θ*_5_) and acceptors (*θ*_3_) are computed. Second (middle column), a statistical model is fitted for each splicing metric that controls for sample covariations and overdispersed count ratios. Third (right column), outliers are detected as data points significantly deviating from the fitted model. Candidates are then visualized with a genome browser.

## Results

To identify splice sites independently of genome annotation, FRASER creates a splice site map by calling *de novo* introns supported by a sufficient amount of RNA-seq split reads (Figure 1, Methods). An intron is defined by a donor (or 5’ splice site) and an acceptor (or 3’ splice site). For each intron, FRASER computes two metrics. The *ψ*_5_ metric quantifies alternative acceptor usage. It is defined as the fraction of split reads from an intron of interest over all split reads sharing the same donor as the intron of interest. The *ψ*_3_ metric, which is analogously defined for the acceptor, quantifies alternative donor usage. FRASER also considers the donor splicing efficiency metric *θ*_5_, defined as the fraction of split reads among split and unsplit reads overlapping a given donor, and the analogously defined acceptor splicing efficiency metric *θ*_3_. Splicing efficiency metrics (collectievly *θ*) have lower values in case of intron retention or impaired splicing. The advantage of these four metrics against alternative splicing metrics such as the popular percent spliced-in^21^ is that they can be quantified from short-read sequencing data without prior exon annotations.^22^ These four metrics are read proportions and therefore range in the interval [0,1]. For modeling and visualization purposes, we used the corresponding log odds ratios that were estimated with a robust logit-transformation (Methods).

To establish FRASER, we considered the Genotype-Tissue Expression (GTEx) project dataset (V7). After quality filtering, this dataset consisted of 7,842 RNA-seq samples from 48 tissues of 543 assumed healthy donors.^20^ Even though the GTEx donors did not suffer from any rare disease, the samples may present aberrant splicing events, just as they do present genes with aberrant expression levels.^23,24^ After filtering for expressed junctions per tissue (Methods), the FRASER splice site map contained in average 137,058 +/- 5,848 donor sites and 136,743 +/- 5,920 acceptor sites (Supplementary Figure S1), where 1.7% and 1.8% of them, respectively, were not in the GENCODE annotation (v28).^25^ Hierarchical clustering of intron-centered logit-transformed *ψ*_5_ values revealed distinct sample clusters for all GTEx tissues (Figure 2A-C). Overall, the average absolute correlation between samples per tissue was 0.10 (+/- 0.05 standard deviation across tissues, Figure 2D). Strong covariation was also observed for *ψ*_3_ and for the splicing efficiency metric (Supplementary Figure S2-3). This covariation structure was tissue-specific (Figure 2A-D). In some tissues, samples clustered according to the RNA degradation index (e.g. heart left ventricle, Figure 2B and Supplementary Figure S2-3), in others to the sequencing center, and in others to the death classification (eg. whole blood samples, Figure 2C). However, no single known covariate could explain covariations for all tissues. Such sample covariations may arise from common genetic variation, technical artefacts or other unknown factors. These observations, consistent with Frésard et al.,^16^ motivated for controlling for between-sample covariations prior calling to aberrant splicing events.

**Figure 2.**
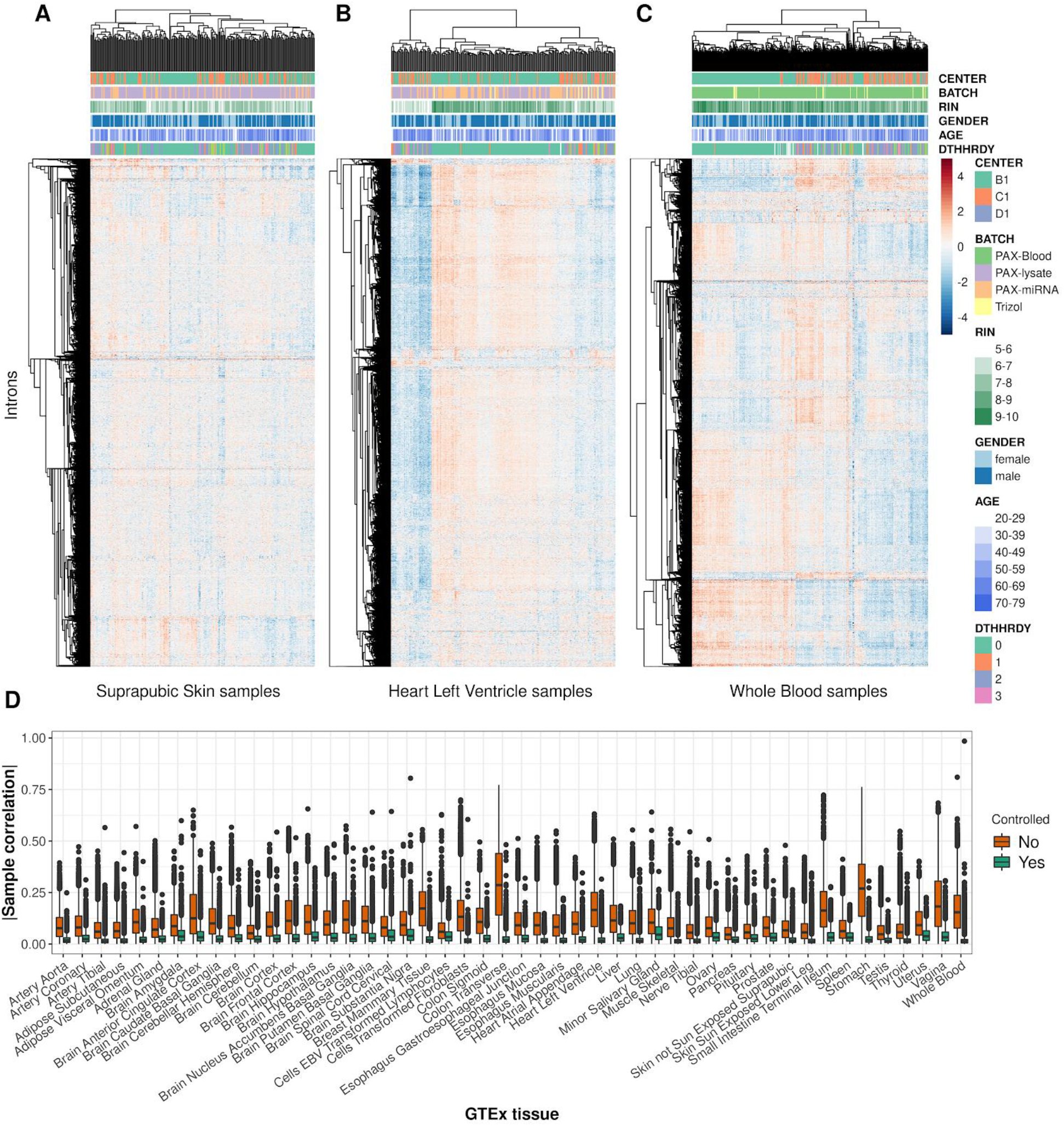
FRASER corrects for covariations in alternative acceptor usage. (**A-C**) Intron-centered and logit-transformed *ψ*_5_ of the 10,000 most variable introns clustered by samples (columns) and introns (rows) for three representative GTEx tissues: suprapubic skin (A), left ventricle heart (B), and whole blood (C). Red and blue depict high and low intron usage, respectively. Colored horizontal tracks display sequencing center, batch, RNA integrity number (RIN), gender, age, and cause of death (DTHHRDY, Hardy scale classification) of the samples. (**D**) Boxplot of absolute values of between-sample correlations of row-centered logit-transformed *ψ*_5_ for 48 GTEx tissues before (orange) and after (green) correction for the latent space. These plots show that while tissue-specific correlation structures exist among samples, latent space fitting allows correcting for them.

We modeled those between-sample covariations by fitting a low-dimensional latent space for each tissue separately. The optimal dimension for the latent space was determined by maximizing the area under the precision-recall curve when calling artificially injected aberrant values independently for each splicing metric (denoising autoencoder, Methods). The method was robust to the exact choice of the encoding dimension, as the performance for recalling artificial outliers typically plateaued around the optimal dimension (Supplementary Figure S4A). The fitted encoding dimension per tissue was 15 for *ψ*_5_, 16 for *ψ*_3_, and 12 for *θ* in average. Moreover, the fitted encoding dimension approximately grew linearly with the number of samples (Supplementary Figure S4B). Larger encoding dimensions were typically found for tissues with more samples (Supplementary Figure S4B). Controlling for the latent space reduced the between-sample correlation from 0.10 +/- 0.05 down to 0.02 +/- 0.01 (mean +/- standard deviation across tissues, Figure 2D and Supplementary Figure S2D and S3D).

### Calling aberrant splicing events using the beta-binomial distribution

Having established an effective procedure to model between-sample covariations, we then addressed the issue of calling aberrant splicing events by finding statistically significant outlier data points. Based on the latent space, FRASER models the expected value of each observation (Methods). We considered the observations that significantly deviate from their expected value as outliers. To this end, we modeled random deviations from the expected values with the beta-binomial distribution, a distribution for count fractions parameterized by its expected count ratio and an intra-class correlation parameter that accounts for variations exceeding sampling noise (Methods). This model allowed computing a two-sided *P* value for each observation (Methods). For the alternative acceptor splicing metric *ψ*_5_, the *P* values of introns with the same donor are not independent since the proportions they are based on sum to one. Therefore, we corrected the *ψ*_5_ *P* values for each donor with the family-wise error rate using Holm’s method, which holds under arbitrary dependence assumption (Methods).^26^ The same was done for *ψ*_3_ yielding a single *P* value per acceptor and sample. We additionally controlled the splice site *P* values for the false discovery rate (FDR) genome-wide per sample using Benjamini-Yekutieli’s method (Methods).^27^ To showcase the application of FRASER, we used the suprapubic skin tissue from the GTEx dataset as done by Brechtmann et al.^28^ Figure 3A shows as an example the *ψ*_5_ metric of the 7^th^ intron of *SRGAP2*, which exhibited a proportional relationship between the number of split reads supporting the 7^th^ intron and the total number of split reads with the same donor site. For this example, the *P* values tended to be conservative yet modeling reasonably well the distribution of the data across samples (Figure 3B). Figure 3C shows as an example with an outlier the *ψ*_5_ metric of the 17^th^ intron of *SRRT*, with one data point exhibiting a much higher usage of this acceptor site than the other samples and a corresponding very low nominal *P* value (*P* = 1.03 x 10^−10^, Figure 3D). Across all introns and splice sites, *P* values were found to be generally conservative (Figure 3E and Supplementary Figure S5). An excess of low *P* values was detected for the most extreme ten thousands of the data, possibly reflecting genuine aberrant splicing events (Figure 3E). Similar results were found for all the investigated GTEx tissues (Supplementary Figure S6).

**Figure 3.**
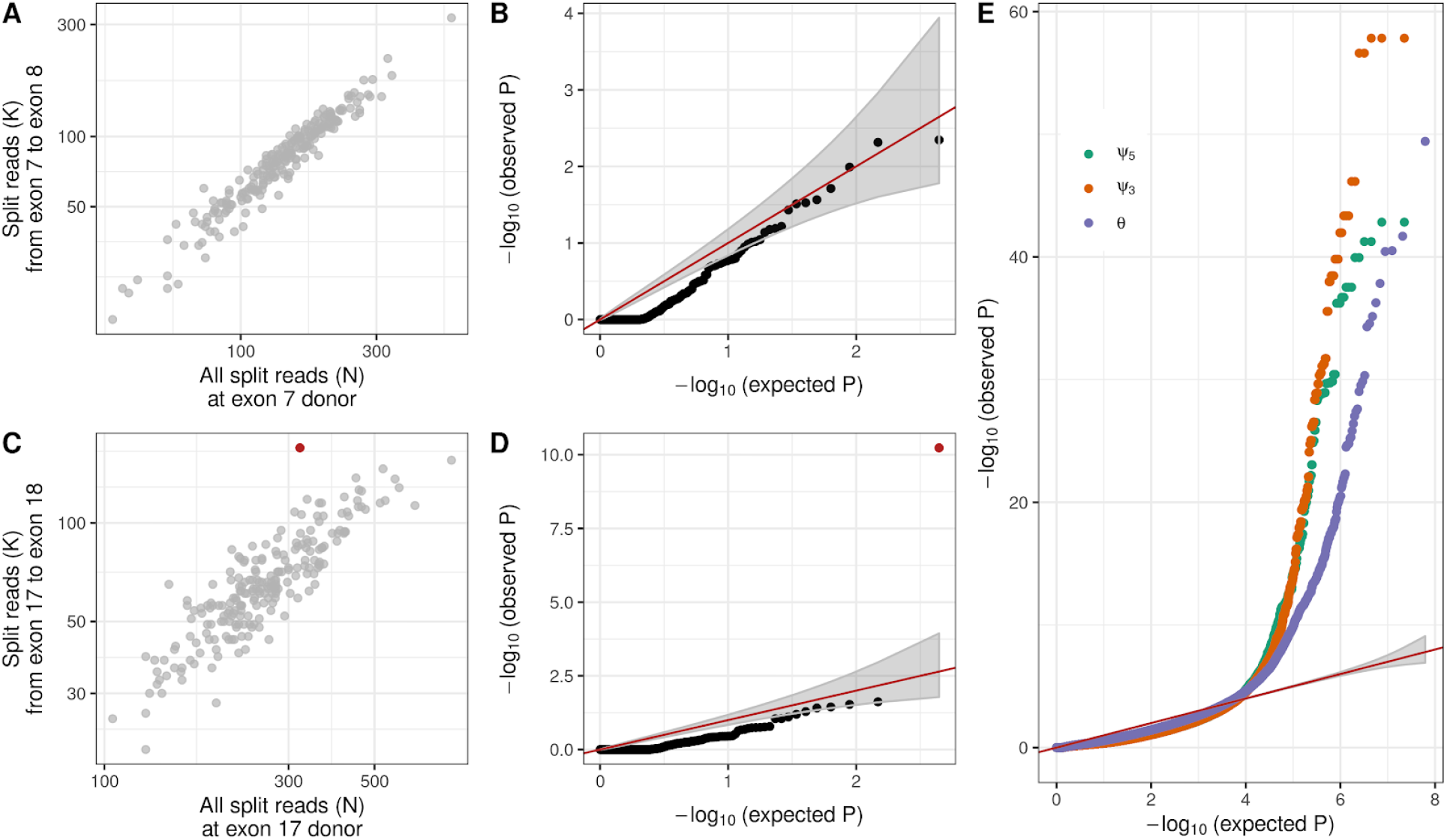
Splicing outlier detection based on the beta-binomial distribution. **(A)** Intron split read counts (y-axis) against the total donor split read coverage for the 7^th^ intron of *SRGAP2*. **(B)** Observed negative log-transformed *P* values (y-axis) against expected ones (x-axis) of the *ψ*_5_ metric for the data shown in A. Under the null hypothesis, the data is expected to lie along the diagonal (red, 95% confidence bands in gray). **(C)** Same as A for the 17^th^ intron of *SRRT*, showing an outlier (FDR < 0.1, red). **(D)** Same as B for the 17^th^ intron of *SRRT*. The outlier is marked in red. **(E)** Same as in B across all introns and splice sites for *ψ*_5_ (orange), *ψ*_3_(green), and splicing efficiency (*θ*, purple). A-E is based on the suprapubic skin tissue from GTEx.

### Recall benchmark of artificially injected outliers

We next aimed at assessing the performance of FRASER and delineating the contribution of modeling the covariation on the one hand, and using the beta-binomial distribution on the other hand. Therefore, we simulated a ground truth dataset based on the suprapubic skin tissue in which we artificially injected splicing outliers with a frequency of 10^−3^, which resulted in 25,988, 26,153 and 49,169 outliers for *ψ*_5_, *ψ*_3_, and *θ*, respectively (Methods). The amplitude of the deviations from the original observed values were drawn uniformly between 0.2 and 1 and their directions (increase or decrease) were randomly assigned with equal probability (Methods). We then monitored outlier recall as well as precision, i.e. the proportion of injected outliers among the reported outliers. Methods not modeling covariation performed worse than methods modeling covariation at any level of recall and for all splicing metrics (Figure 4 and Supplementary Figure S7-8). Moreover, methods modeling covariation and using beta-binomial based *P* values yielded a higher precision than those using z scores (Figure 4 and Supplementary Figure S7-8). This higher precision was observed at all levels of recalls, simulated outlier amplitudes, and read coverage (Figure 4). Notably, using PCA and a z score cutoff equal to 2 instead of FRASER at FDR < 0.1 yielded two orders of magnitude more outliers across all GTEx tissues (Supplementary Figure S9) and a drastic drop in precision (3% versus 92% with FRASER) for a small increase in recall (98% versus 83% with FRASER, Supplementary Figure S10). This drastic difference of precision strongly suggests using an FDR cutoff rather than an absolute z score cutoff.

**Figure 4.**
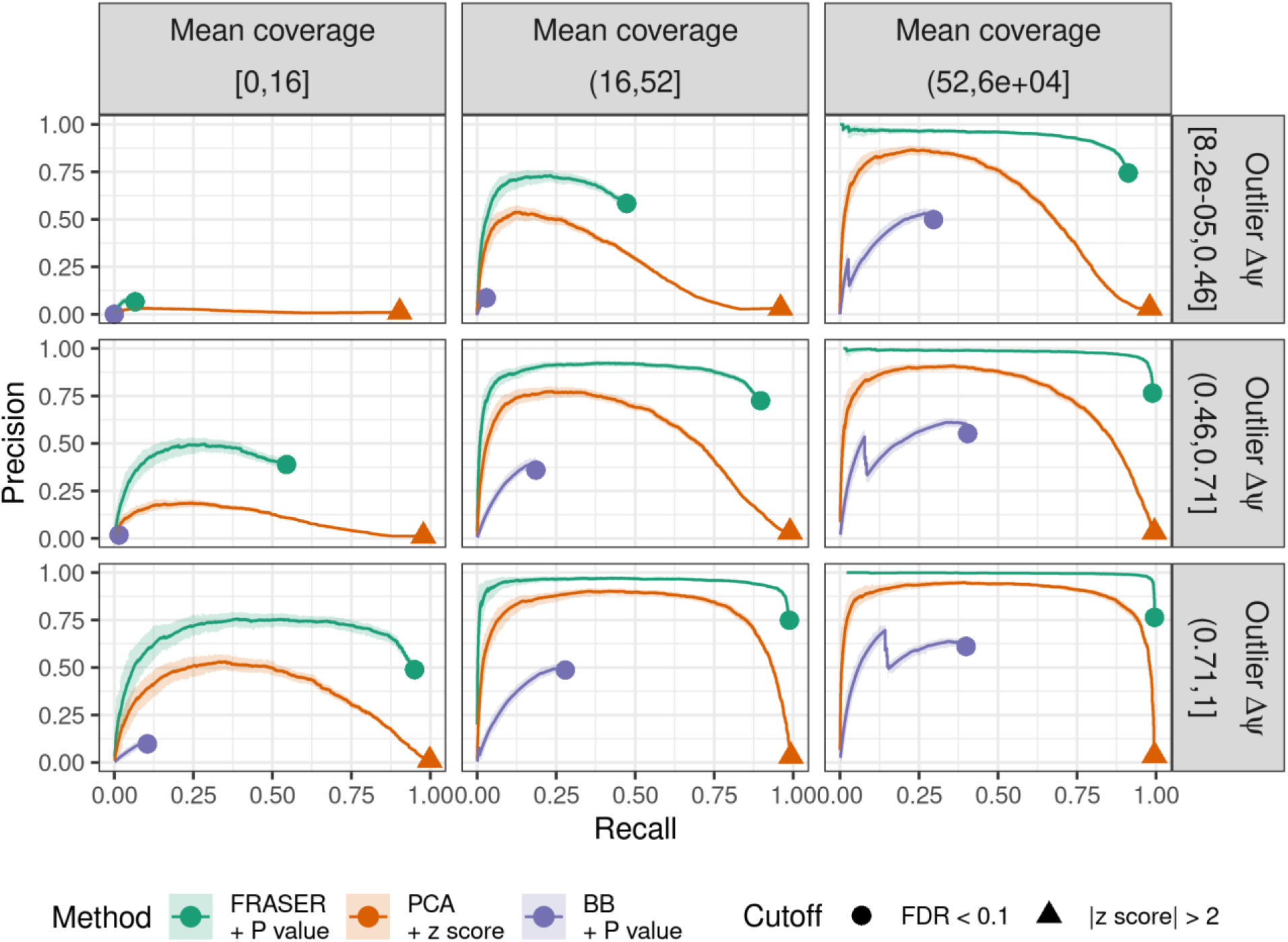
Benchmark using artificially injected outliers for alternative acceptor usage. The proportion of simulated outliers among reported outliers (precision) plotted against the proportion of reported simulated outliers among all simulated outliers (recall) for increasing *P* values (FRASER, green and beta binomial, purple) or decreasing absolute z scores (PCA, orange). Additionally, all events with |*Δψ*_5_| < 0.1 are ranked last. The data is stratified by the mean coverage of the intron (columns) and by the injected absolute *Δψ_5_* value (rows). The cutoffs for each method are marked (FDR < 0.1, circle, and absolute z score > 2, triangle). Light ribbons around the curves depict 95% confidence bands estimated by bootstrapping. These results show the importance of controlling for latent confounders, using a count-based distribution, and correcting for multiple testing.

The benchmark with simulated outliers also allowed investigating alternative ways to estimate the expected values by regression on the latent space. This included a beta-binomial regression, a robust version of the beta-binomial regression, as well as a least squares regression of logit-transformed splicing metrics (Methods). The latter approach, which we eventually adopted for FRASER, had similarly high performance as the robust beta-binomial regression (Supplementary Figure S7-8, Methods) while being much faster to compute. The beta-binomial regression was too sensitive to outlier data points and hence was outperformed by its robust version (Supplementary Figure S11).

### Rare variant enrichment analysis

We further evaluated the performance of FRASER by assessing the enrichment of rare genetic variants among splicing outlier genes with the rationale that some aberrant splicing events are caused by rare genetic variants. For this analysis, we defined a variant to be rare when having a minor allele frequency (MAF) less than 0.05 within GTEx^20^ and gnomAD^29^ as done for gene expression in Li et al.^23^ We annotated these variants in two ways. First, we considered splice region variants (Methods), which we defined as variants located within 1-3 bases of an exon or 1-8 bases of an intron. This correspond to the union of the splice site dinucleotide and splice region variants as defined by the sequence ontology through the variant effect predictor (VEP).^30,31^ We found on average 299.4 +/- 207.6 rare splice region variants. Second, we considered rare variants predicted to affect splicing by MMSplice,^8^ a machine learning algorithm that scores variants as far as 100 base pairs away from splice sites (on average 66.0 +/- 48.0 rare MMSplice variants, Methods). The consequences of a genetic variant on splicing may spread across splice sites of a gene, due to complex effects including competition between splice sites, or coordinated splicing between distant exons.^32^ Hence, the detectable effects of a variant affecting splicing may not necessarily be at its closest splice sites. Therefore, we computed the enrichment at the gene level. To this end, we computed gene-level *P* values using a family wise error rate correction across all splice sites within a gene (Methods). Additionally to the previously benchmarked methods, we also applied the method of Kremer et al.,^15^ which is based on the gene-level differential splicing algorithm Leafcutter.^18^

Across all 48 GTEx tissues, FRASER showed higher enrichments than Leafcutter, Gaussian-based *P* values, and non-corrected beta-binomial *P* values. The higher enrichments observed held for different nominal *P* value cutoffs and both for rare variants in the splice regions, as well as for those predicted to affect splicing by MMSplice (Figure 5 and Supplementary Figure S12). Notably, the MMSplice variant set showed 2 to 10 times higher enrichments across all methods compared to the splice region variant set, emphasizing the importance of considering exonic or deep intronic variants as potential splice-affecting candidates. Altogether, this gene-level benchmark on non-simulated data confirms the importance of controlling for covariation and using a count fraction distribution to identify aberrant splicing. This benchmark also shows that FRASER outperforms both state-of-the-art methods, which are the Leafcutter based approach^15^ and the z score based approach.^16^

**Figure 5.**
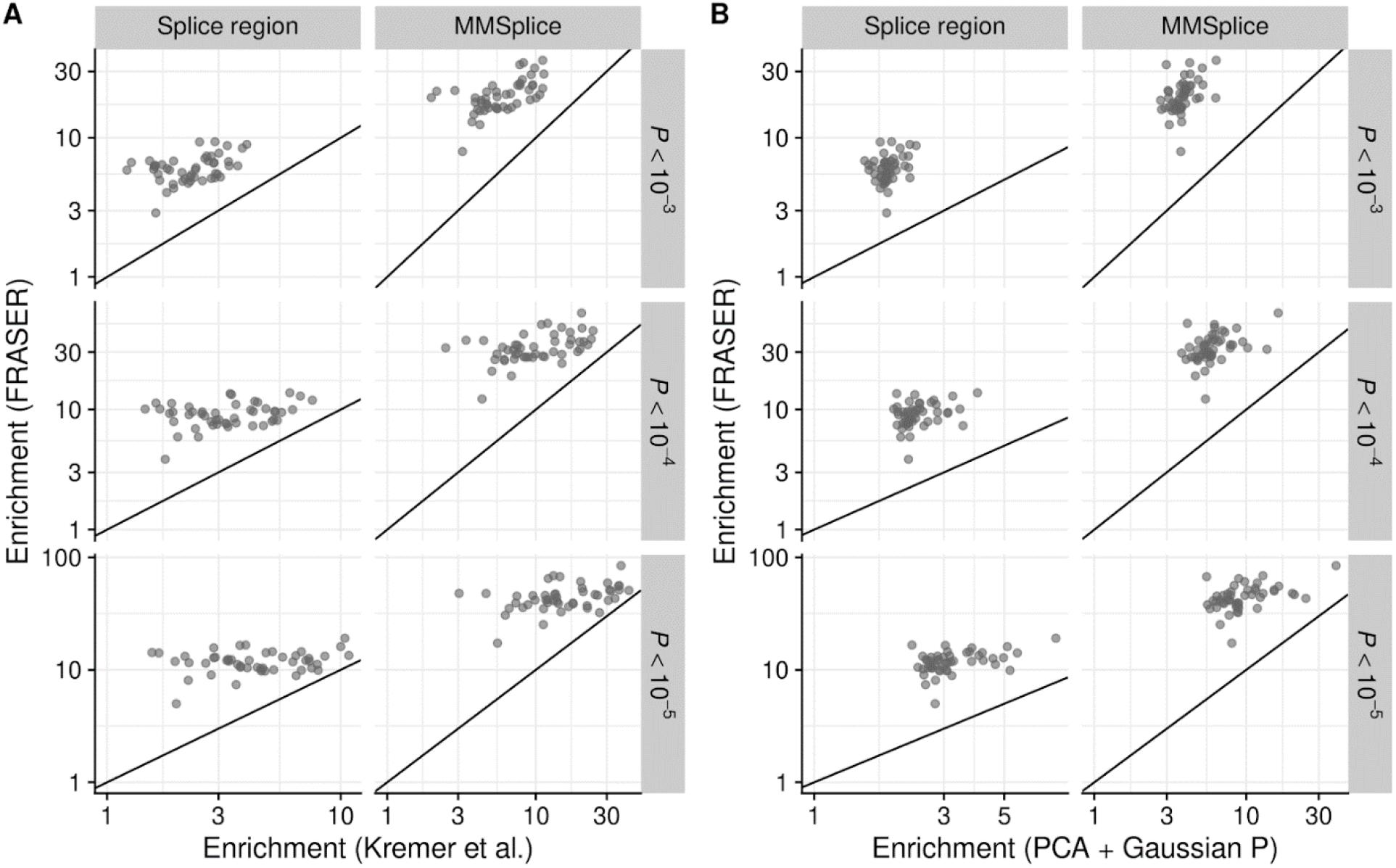
Enrichment for rare variants. **(A)** Enrichment using FRASER (y-axis) against enrichment using the splicing detection approach from Kremer et al.^15^ (x-axis) for rare variants located in a splice region or for rare variants predicted to affect splicing by MMSplice^8^ (column). The enrichment is calculated for different nominal *P* value cutoffs (rows). Each dot represents a GTEx tissue. **(B)** The same as A but the enrichment of FRASER (y-axis) is compared against Gaussian based *P* values on top of PCA controlled splicing metrics (x-axis). In every panel, the enrichment for each of the 48 GTEx tissues is equal or higher for FRASER (points above the diagonal line).

### Application to rare disease diagnosis

Having established FRASER using a large cohort of healthy donors, we next reanalyzed the 119 RNA-seq samples from skin fibroblasts of 105 individuals with a suspected rare mitochondrial disorder of Kremer et al. (hereafter the Kremer dataset).^15^ In a rare disease diagnosis context, the aim is to identify aberrant splicing events that could be disease causing, typically by disrupting the function of a phenotypically relevant gene. To this end, gene-level statistics are handier entry points than splice-site level statistics. Moreover, we suggest combining statistical significance cutoffs with effect size cutoffs because larger effects are more likely to have strong physiological impacts. For the Kremer dataset, FRASER reported a median of 12, 7, and 10 genes with at least one aberrant splicing event per sample for *ψ*_5_, *ψ*_3_, and splicing efficiency, respectively, at a significance level of FDR < 0.1 and an effect size greater than 0.3 (absolute difference between observed and expected value, Figure 6A). Similar amounts were found on all 48 GTEx tissues (Supplementary Figure S9). These criteria yielded slightly less splicing outliers than in the original study (1666 versus 1725, Figure 6B) yet detecting all novel pathogenic splice events reported in the original study (*CLPP*, *TIMMDC1* in 2 individuals, and *MCOLN1*, Figure 6B). Notably, the intron retention event in the gene *MCOLN1* was missed by the aberrant splicing pipeline used by Kremer et al.^15^ because it was based on Leafcutter which does not consider non-split reads. (Kremer et al. identified the event through mono-allelic expression of the heterozygous intronic variant.) Generally, including the splicing efficiency metrics with FRASER led to a two-fold increase of detected aberrant events over considering the alternative splicing metrics *ψ*_5_, and *ψ*_3_ alone (Figure S13). Altogether, these findings show the clinical relevance and the complementarity of using both splicing efficiency and alternative splicing metrics.

**Figure 6.**
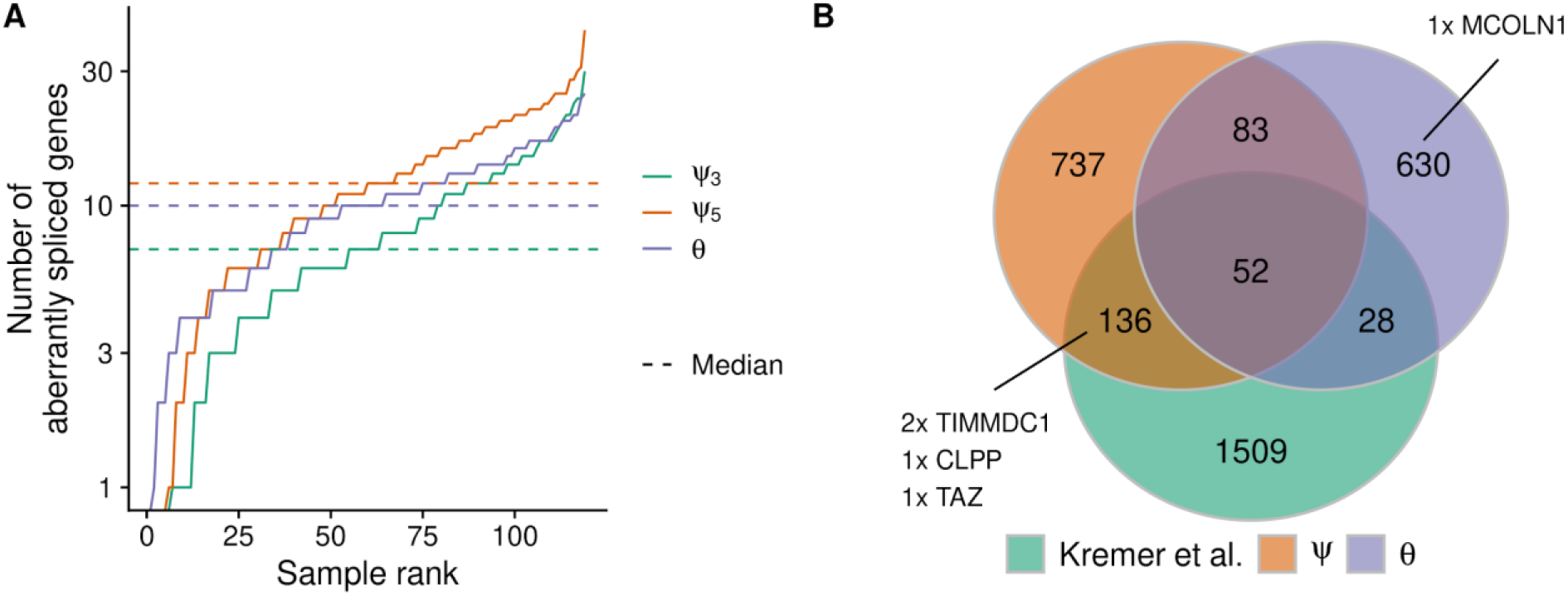
Aberrant splicing detection in a rare disease cohort. **(A)** Number of aberrantly spliced genes within the Kremer dataset (FDR < 0.1 and |*Δψ*| > 0.3) per sample ranked by the number of events for *ψ*_5_ (orange), *ψ*_3_ (green), and *θ* (purple). **(B)** Venn diagram of the aberrant splicing events detected by FRASER using alternative splicing (orange, *ψ*) or splicing efficiency (violet, *θ*) only and by Kremer et al. (green).^15^ Pathogenic splicing events are labelled with the gene name.

Moreover, the reanalysis of the rare disease dataset highlighted aberrant alternative donor usage in the gene *TAZ* for the undiagnosed individual 74116 (difference *ψ*_3_ = −0.88 and FDR = 1.98 x 10^−9^, Figure 7A), which was overlooked in the original study.^15^ The nearly complete loss of the canonical donor site usage of the 4^th^ exon (Figure 7B-D) resulted in the usage of a newly created donor site located 22 bp inside the 4^th^ exon (Figure 7E). Usage of the new donor site leads to an ablation of 8 amino acids of the protein encoded by *TAZ*, Tafazzin. Tafazzin catalyzes maturation of cardiolipin, a major lipid constituent of the inner mitochondrial membrane involved in energy production and mitochondrial shape maintenance.^33^ Moreover, individual 74116 harbors a rare homozygous synonymous variant (c.348C>T) that creates the new upstream donor site by introducing a GT dinucleotide (Figure 7E). This variant had not been prioritized by WES analysis as it was synonymous and not indexed by ClinVar.^34^ However, the variant had been previously associated with a splicing defect in *TAZ* and dilated cardiomyopathy,^35^ consistent with individual 74116’s myopathic facies and arrhythmias, thereby establishing the genetic diagnosis.

**Figure 7.**
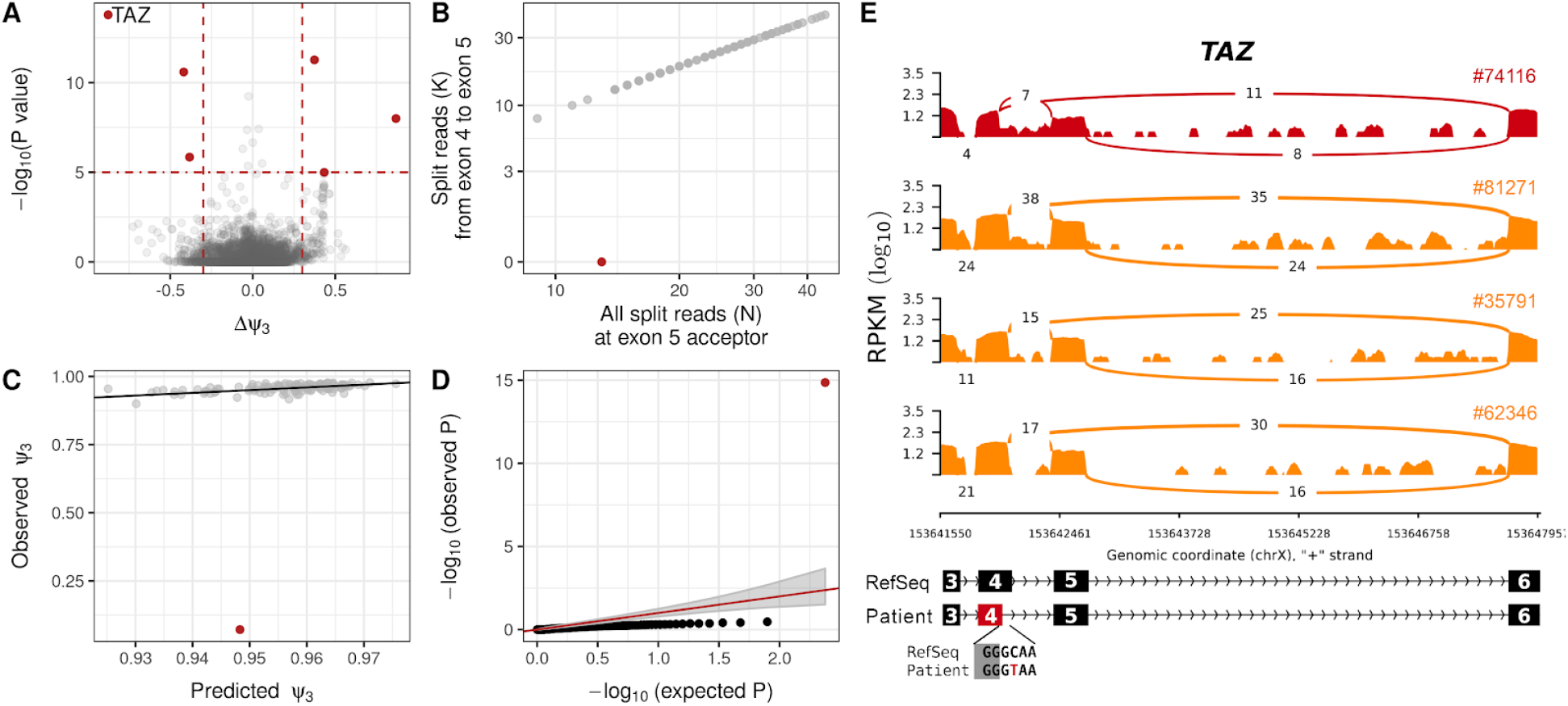
Detection of a pathogenic splicing defect using FRASER. **(A)** Gene-level significance (−log_10_(*P*), y-axis) versus effect (observed minus expected *ψ*_3_, x-axis) for alternative donor usage for the individual 74116. Five genes (red dots, among them *TAZ*) passed both the genome-wide significance cutoff (horizontal dotted line) and the effect size cutoff (vertical dotted lines). **(B)** Number of split reads spanning from the 4^th^ exon to the 5^th^ exon (y-axis) against the total number of split reads at the acceptor site of the 5^th^ exon (x-axis) of the *TAZ* gene. Sample 74116 (red) deviates from the cohort trend. **(C)** Observed (y-axis) against FRASER-predicted (x-axis) *ψ*_3_ values for data in B. **(D)** Quantile-quantile plot of observed *P* values (−log_10_(*P*), y-axis) against expected *P* values (−log_10_(*P*), x-axis) and 95% confidence band (gray) for data in B. Sample 74116 (red) shows an unexpected low *P* value. **(E)** Sashimi plot of the exon-truncation event in RNA-seq samples of the *TAZ*-affected (red) and three representative *TAZ*-unaffected (orange) individuals. The RNA-seq read coverage is given as the log_10_ RPKM-value (Reads Per Kilobase of transcript per Million mapped reads, y-axis) and the number of split reads spanning an intron is indicated on the exon-connecting line. At the bottom, the gene model of the RefSeq annotation is depicted in black and the aberrantly spliced exon is coloured in red. The insert depicts the donor-site creating variant of the affected individual 74116.

### Implementation

FRASER is implemented as an R/Bioconductor package.^36,37^ It contains functions to count RNA-seq reads, fit the model, calculate *P* values, as well as to extract and visualize the results. The workflow and functionalities of the FRASER package are aligned with the previously published OUTRIDER package.^28^ The package allows for a full analysis to be made with only a few lines of code and comes along with a comprehensive vignette guiding through a typical analysis. It is available as open source through https://github.com/gagneurlab/FRASER.

## Discussion

We have introduced FRASER, an algorithm specifically developed for detecting aberrant splicing events in RNA-seq data. The combination of three features make FRASER unique: (i) it considers non-split reads overlapping splice sites, allowing for detecting intron retention, (ii) it automatically controls for latent confounders, and (iii) it assesses statistical significance with a count distribution. Extensive benchmarks with artificially simulated aberrant splicing events, enrichment of rare variants with a splicing effect potential, as well as reanalysis of a rare disease cohort demonstrated the importance of each of these features. FRASER is provided as an easy-to-use R/Bioconductor package.

We implemented FRASER in a modular way so that the procedures for fitting the latent space and for estimating expected values given the latent space can be independently chosen as well as the distributions used to define splicing outliers. The best performing model was obtained using a hybrid combination in which fitting the latent space and estimating expected values are performed with a least-squared loss while the beta-binomial distribution is used for assessing the significance of the outlier. This combination does not correspond to a maximum likelihood fit of a particular distribution we are aware of, but it gave the best empirical results. Future research could investigate whether some other classes of models such as a multivariate logit-normal binomial distribution could provide good maximum likelihood fits to splicing metrics.

FRASER is based on splicing metrics defined at the level of individual splice sites. In theory, using a gene model that integrates data across entire splice isoforms can increase sensitivity because all reads supporting an isoform over another contribute to the test statistic. However, the difficulty is that either a gene model must be known beforehand or it has to be assembled de novo. While we were developing FRASER, a preprint described a splicing outlier detection method that fits a Dirichlet-Multinomial distribution at the gene level on all split read counts of alternatively spliced exon-exon junctions called de novo.^24^ The method appears to require robust gene models in the first place, because data preprocessing and filters yielded only 6,000 genes analysed on a typical GTEx tissue. This filtering and clustering may be appropriate to investigate healthy populations and basic biology of aberrant splicing,^24^ but limits the usability in rare disease diagnostics, where a single event could be the disease-causing one.

One limitation of RNA-seq for diagnosis of rare diseases is that the affected tissue may not be accessible. Nonetheless, a causal splicing defect may also be detectable in a clinically accessible tissue such as blood or skin, while its pathological consequence may be revealed only in the affected tissue. The *TAZ* gene is such an example with pathological effects in heart but with normal gene expression levels in skin despite the splice defect. We suggest investigators to check the gene and exon overlap with tissues of interest, using the MAJIQ-CAT web interface^38^ or to refer to the full splice site map of the GTEx tissues we have compiled here (Web Resources). Moreover, this GTEx splice site map can be used to extend RNA-seq compendia for users with only a few samples. In conclusion, through the easy to use R/Bioconductor package and the ability to utilize the preprocessed GTEx dataset, we foresee FRASER to become an important tool for the growing field of RNA-seq based diagnosis of rare diseases.

## Methods

### Datasets

We considered two RNA-seq datasets: (i) a dataset consisting of 119 RNA-seq samples from skin fibroblasts of 105 individuals with a suspected rare mitochondrial disease^15^ (the Kremer dataset) and (ii) 7,842 RNA-seq samples from 48 tissues of 543 assumed healthy individuals of the Genotype Tissue Expression Project V7^20^ (hereafter the GTEx dataset). Both datasets are not strand-specific. Read mapping files in the BAM file format were obtained for the Kremer dataset by mapping the RNA-Seq reads to the UCSC hg19 genome assembly^39^ with the STAR aligner (version 2.4.2a).^40^ To detect novel exon junctions, we ran STAR in the two-pass mode (option *twopassMode=Basic*) with minimal chimeric segment length of 20 (*chimSegmentMin=20*). For GTEx, we obtained the BAM files from dbGaP (phs000424.v7.p2), which were already aligned with STAR (version 2.4.2a). The GTEx consortia used the same parameters except mapping against the GRCh37 genome assembly based on the GENCODE v19 annotation.^25^ We considered only samples with an RNA integrity number of 5.7 or higher and marked as usable by the GTEx consortia (SMRIN and SMAFRZE column, respectively) and discarded tissues with less than 50 samples.

### Read Counting and splicing metrics

The set of acceptor and donor splice sites (or the splice site map) of a dataset was defined by calling all introns including *de novo* events based on RNA-seq split reads. To this end, the split reads were extracted from the BAM files and counted using the R/Bioconductor packages GenomicAlignments and GenomicRanges.^41^ Having defined the splice site map, non-split reads overlapping splice sites were counted in order to compute the splicing efficiency, which can be used to detect intron retention. Specifically, the non-split reads were counted for each splice site using the R/Bioconductor Rsubread package^42^ requiring at least 5 nt aligned on each side of the splice site to be robust against mapping errors of very short overhangs as done in Braunschweig et al.^43^

As described by Pervouchin et al.,^22^ we compute for each sample, for donor *D* (5’ splice site) and acceptor *A* (3’ splice site) the *ψ*_5_ and *ψ*_3_ values, respectively, as:

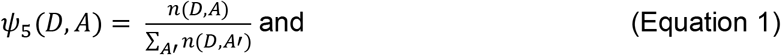

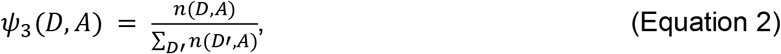

where *n*(*D, A*) denotes the number of split reads spanning the intron between donor *D* and acceptor *A* and the summands in the denominators are computed over all acceptors found to splice with the donor of interest (Equation 1), and all donors found to splice with the acceptor of interest (Equation 2). To not only detect alternative splicing but also partial or full intron retention, we considered a splicing efficiency metric. Multiple related definitions exist including 3’ SS ratio,^44^ completeness of splicing index,^45^ and percent intron retained.^43^ We used the *θ*_5_ and *θ*_3_ values as defined by Pervouchin et al.^22^ Specifically:

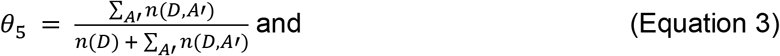

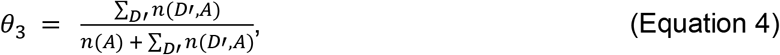

where *n*(*D*) is the number of non-split reads spanning exon-intron boundary of donor *D*, and *n*(*A*) is defined as the number of non-split reads spanning intron-exon boundary of acceptor *A*. While we calculate *θ* for the 5’ and 3’ splice sites separately, we do not distinguish later in the modeling step between *θ*_5_ and *θ*_3_ and hence call it jointly *θ*.

For robust fitting of the model, we restricted the analysis to splice sites of introns supported by at least 20 split reads in at least one sample. Further, we filtered out splice sites and introns where more than 95% of the samples had zero coverage.

### Statistical Model

The metrics *ψ*_5_, *ψ*_3_, and *θ* are count proportions. For each of these metrics, we model the distribution of the numerator conditioned on the value of the denominator using the beta-binomial distribution. Unlike the binomial distribution, the beta-binomial distribution can account for overdispersion. Specifically, for *ψ*_5_, we assume that the split read count *k_ij_* of the intron *j* = 1,…, *p* in sample *i* = 1,…, *N* follows a beta-binomial distribution with an intron-specific intra-class correlation parameter *ρ_j_* and a sample- and intron-specific proportion expectation *μ_ij_*:

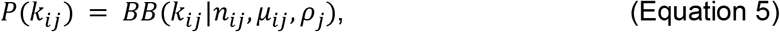

where *n_ij_* defines the total number of split reads having the same donor site than intron *j*. The metrics *ψ*_3_ and *θ* are modeled analogously. For ease of writing we will refer in the following with *ψ* always to the site-specific *ψ*_5_ and *ψ*_3_ form. Both *μ_ij_* and *ρ_j_* are limited to the range [0,1]. The parametrization of the beta-binomial distribution we used can be found in the Supplemental Material and Method section.

The proportion expectation *μ_ij_* is jointly modeled using a latent space that captures covariations between samples. Specifically, we model:

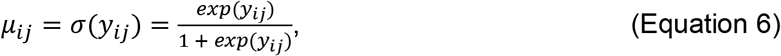

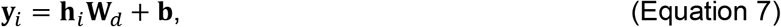

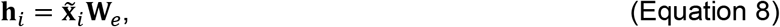

where the vectors **h**_*i*_ are the rows of the matrix **H**, the *N* × *q* projection of the data onto the *q*-dimensional latent space with 1 < *q* < min(*p, N*), **W**_*e*_ is the *p* × *q* encoding matrix, **W**_*d*_ the *q* × *p* decoding matrix, and the *p*-vector **b** is a bias term. The input vector 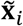 is given by the centered and logit-transformed pseudo-count ratios. We define 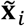 as:

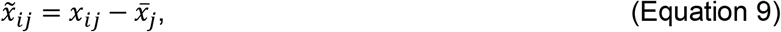

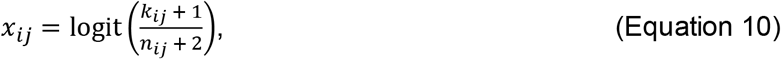

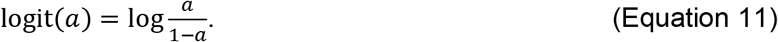

### Fitting of the latent space and the distribution

Four parameters must be fitted, namely **W**_*e*_, the encoding matrix, **W**_*d*_, the decoding matrix, **b**, the bias term, and *ρ_j_*, the intra-class correlation of the beta-binomial distribution. The fitting of these parameters is achieved in two steps. First, the latent space **H** and the expected splicing proportions *μ_ij_* are fitted with a principal component analysis (PCA). To this end, a PCA is computed on the input matrix 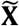 using the pcaMethod package.^46^ The latent space **H** is then computed with Equation (8) by setting the encoder matrix **W**_*e*_ to the first *q* loadings of the PCA. Given the latent space **H**, *μ_ij_* is computed by using the transpose of **W**_*e*_ for **W**_*d*_ and setting the bias term to 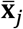. In the second step, the intra-class correlation parameter *ρ_j_* is fitted given the count proportion expectations using a beta-binomial loss function. Specifically, we use the *optimize* function from R^37^ and minimize the average negative beta-binomial log-likelihood in parallel across introns (Supplemental Material and Method).

Additionally, we implemented an alternative approach to fit the distribution parameters given the latent space **H**. To this end, we use a weighted negative beta-binomial log-likelihood loss function to model in an iterative fashion the decoding matrix **W**_*d*_ and the bias term **b** on the one hand and the intra-class correlation parameter *ρ* on the other hand. First, we initialize the parameters as described before using PCA. Given the latent space, these parameters can be fitted independently for each intron *j*. We start by optimizing *ρ_j_* given the weights 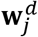 and the bias *b_j_* (step 1). Then, we optimize 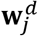 and *b_j_* given *ρ_j_* in step 2. Steps 1 and 2 are repeated until the average weighted negative log-likelihood of each step in one iteration does not differ more than the convergence threshold of 10^−5^ from the last step of the previous iteration, or 15 iterations are reached, which triggers a warning. We use the L-BFGS method implemented in the R function *optim* to fit the decoder weights and the bias.^47^ A detailed derivation of the used loss functions and the respective gradients can be found in the Supplemental Material and Method section.

### Robust distribution fitting using a weighted log-likelihood

Outlier data points can have strong effects on the model fitting. We use weights in the loss function to decrease the influence of outliers on the model, following the edgeR approach.^48^ Specifically, we define the weight for each observation based on its Pearson residual. The Pearson residual of the observed data point *x_ij_* (Equation 10) with respect to the beta-binomial distribution including the pseudocounts is defined as follows:

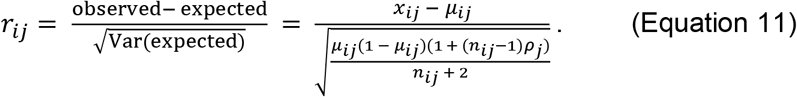

The weights *w_ij_* for sample *i* and intron *j* are obtained from these residuals using the Huber function:^42^

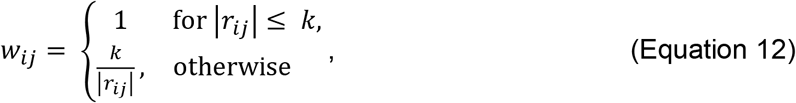

where we use *k* = 1.345 as suggested in the edgeR package,^48^ which leads to downweighting of about 5% of the data points. These weights are then included in the calculation of the log-likelihood yielding the average weighted log-likelihood *L^W^*:

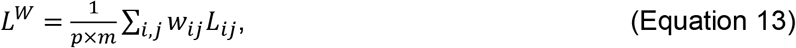

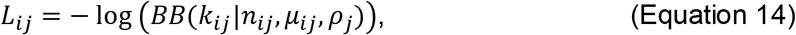

where *L_ij_* is the beta-binomial log-likelihood of sample *i* and intron *j* as defined in the Supplemental Material and Method section.

### Finding the Hyperparameters

The model fitting procedure described above leaves one hyperparameter that remained to be optimized: the latent space dimension *q*. In order to find the optimal latent space dimension *q*, we implemented a denoising autoencoder approach.^19^ Specifically, we generated corrupted data by injecting aberrant read count ratios with a frequency of 10^−2^ into the original data. The injection scheme is laid out in detail in the next section. We then select *q* as the dimension maximizing the area under the precision-recall curve for identifying corrupted read ratios. This is done for each splicing metric separately. To speed up the fitting procedure of the hyperparamtere, we randomly subset the input matrix 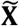 to 15,000 introns out of the 30,000 most variable introns with a mean total coverage greater than 5. This subsetting is done before the injection of aberrant read count ratios.

### Injection of artificial outliers

To fit the FRASER hyperparameter as well as to compare the outlier detection performance between FRASER and other methods, we developed a procedure to inject artificial outliers into a given dataset. For injection, we considered all expressed introns or splice sites within the dataset and injected only one outlier per splice site and sample. Outliers were randomly injected with a frequency of 10^−2^ for the hyperparameter optimization and with a frequency of 10^−3^ for the benchmarking.

To create aberrant splicing ratios, we inject *in silico* a splicing outlier count *k_ij_^o^* by changing the original read count *k_ij_* such that the value of changes by *Δψ_ij_^o^*. *Δψ_ij_^o^* is drawn from a uniform distribution:

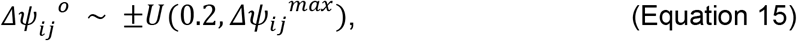

where *Δψ_ij_^max^* is the maximal possible *Δψ_ij_* for intron *j* in sample *i*. The value of *Δψ_ij_^max^* is dependent on the randomly sampled injection direction: *Δψ_ij_^max^* = 1 − *ψ_ij_* and *Δψ_ij_^max^* = *ψ_ij_* for up- or downregulation, respectively. To ensure that an aberrant splice ratio can be injected the direction is switched if *Δψ_ij_^max^* < 0.2. We injected outliers only for introns harboring 10 reads or more in the considered sample.

Taking the pseudocounts into account, the outlier count *k_ij_^o^* is then given by

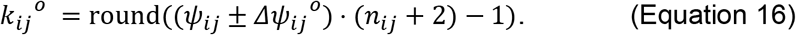

In order to provide a biologically realistic outlier injection scheme that preserves the total amount of reads, the counts for the introns *l* sharing the same donor or acceptor, respectively, with *k_ij_^o^* are changed accordingly, where the *Δψ_ij_^o^* change is distributed equally over all secondary introns *l* as follows:

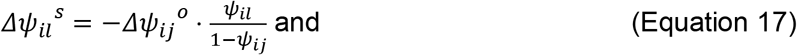

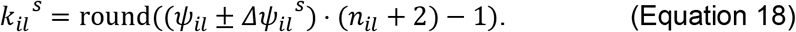

### Statistical significance

Statistical significance of outliers is performed by testing the null hypothesis that the count *k_ij_* with *n_ij_* trials follows a beta-binomial distribution with parameters fitted as described above for every pair of sample *i* and intron *j*. We compute two-sided *P* values *p_ij_* using the mean probability of success *μ_ij_* and the fitted intra-class correlation parameter *ρ_j_* as follows:

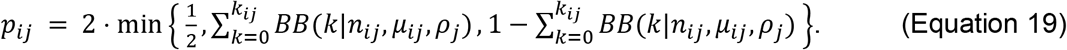

The term ½ is included to prevent getting *P* values greater than 1, which can happen due to the nature of the discrete distributions.

The *P* values of introns sharing a splice site are not independent since the proportions they are based on sum to one. We therefore correct the *P* values for each splice site with the family-wise error rate (FWER) using Holm’s method, which holds under arbitrary dependence assumption,^26^ and report the minimal corrected *P* value per splice site. An additional FWER step is performed on the gene level if gene-level *P* values are requested. To correct for multiple testing genome-wide, we use the Benjamini-Yekutieli false discovery rate (FDR) method^27^ as both splice site corrected *P* values and the gene-wise corrected *P* values can still be correlated due to biological effects that are not completely removed by the model. All *P* value corrections are done per sample.

### Z Score and *Δψ* Calculation

Z scores *z_ij_* are calculated per intron on the difference on the logit scale between the measured *ψ* value including pseudocounts and the proportion expectation *μ_ij_* as:

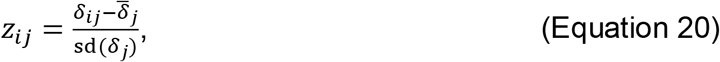

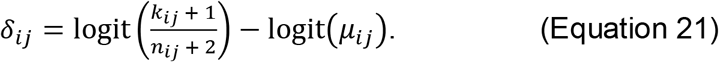

The *Δψ* values are calculated as the difference between the observed *ψ_ij_* value on the natural scale including pseudocounts and the proportion expectations *μ_ij_*:

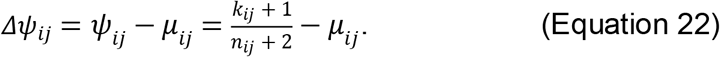

### Alternative Splicing Outlier Detection Methods

We implemented different alternative splicing outlier detection methods to assess the performance of FRASER. As the baseline for our approach, we used a simple beta-binomial distribution (BB) with no correction for existing covariation and the parameters *μ_ij_* and *ρ_j_* were estimated with the R package VGAM.^49^ Further, we implemented a Z score approach similar to the approach described by Frésard et al.^16^ Instead of regressing out the top *q* principal components accounting for 95% of the variation within the data, we used the top *q* loadings of the PCA maximizing the precision-recall of *in silico* injected splicing outliers and computed the Z scores according to Equation (16). Finally, we implemented the Leafcutter^18^ approach described by Kremer et al,^15^ where one sample is compared against all others within the dataset and no control for latent sources of sample covariation is considered.

### Enrichment analysis

For the GTEx enrichment analysis, we obtained all rare variants (MAF < 0.05 within all 635 GTEx samples and in gnomAD^29^) from the GTEx whole genome sequencing genetic variant data (V7). From this rare variant set, we extracted on the one hand all annotated splicing variants (*splice_donor*, *splice_acceptor*, and *splice_region*) according to the sequencing ontology and VEP.^30,31^ This covers all variants around the exon-intron and intron-exon boundary, which is 1-3 bases within the exon and 1-8 bases within the intron. On the other hand, we extracted variants predicted to affect splicing by MMsplice.^8^ To this end, we scored all variants within 100 bp of an annotated exon (GENCODE v30)^25^ with MMSplice in an exon-centric way. Then, variant-exon pairs with a score of |*Δ*logit(*ψ*)| > 2 were selected. We then computed enrichments for rare splicing variants found within outlier genes as the proportion of outliers having a rare splicing variant over the proportion of non-outliers having a rare splicing variant.

## Description of Supplemental Data

The Supplemental Data includes 13 figures. The Supplemental Material and Methods section includes detailed derivations of the used loss functions and their respective gradients.

## Acknowledgments

C.M., V.A.Y., H.P., and J.G. were supported by the EU Horizon2020 Collaborative Research Project SOUND (633974). The Bavaria California Technology Center supported C.M through a fellowship. The German Bundesministerium für Bildung und Forschung (BMBF) supported the study through the e:Med Networking fonds AbCD-Net (FKZ 01ZX1706A to V.A.Y., C.M., and J.G.), the German Network for Mitochondrial Disorders (mitoNET; 01GM1113C to H.P.), and the E-Rare project GENOMIT (01GM1207 to H.P.). A fellowship through the Graduate School of Quantitative Biosciences Munich supports V.A.Y. The Genotype-Tissue Expression (GTEx) Project was supported by the Common Fund of the Office of the Director of the National Institutes of Health and by the National Cancer Institute, National Human Genome Research Institute, National Heart, Lung, and Blood Institute, National Institute on Drug Abuse, National Institute of Mental Health, and National Institute of Neurological Disorders and Stroke. The data used for the analyses described in this manuscript were obtained from the GTEx Portal on April 4, 2019, under accession number dbGaP: phs000424.v7.p2.

## Author contributions

C.M. and J.G conceived the method. C.M and I.S implemented the package and performed the full analysis. V.A.Y. contributed to the package development and to the analysis. M.H.Ç. performed the MMSplice analysis of GTEx. C.M. and Y.L. performed the rare variant enrichment analysis. L.S.K. and M.G. analyzed the results of the rare disease cohort. J.G and H.P. supervised the research. C.M., I.S, and J.G. wrote the manuscript with the help of V.A.Y. All authors revised the manuscript.

## Web Resources

GTEx Portal, https://www.gtexportal.org/home

FRASER package, http://github.com/gagneurlab/FRASER

FRASER analysis pipeline, https://github.com/gagneurlab/FRASER-analysis

GTEx splice map, https://i12g-gagneurweb.in.tum.de/public/paper/FRASER

## Reference

1. López-Bigas, N., Audit, B., Ouzounis, C., Parra, G. & Guigó, R. Are splicing mutations the most frequent cause of hereditary disease? FEBS Lett. 579, 1900–1903 (2005).

2. Wang, G.-S. & Cooper, T. A. Splicing in disease: disruption of the splicing code and the decoding machinery. Nat. Rev. Genet. 8, 749–761 (2007).

3. Park, E., Pan, Z., Zhang, Z., Lin, L. & Xing, Y. The Expanding Landscape of Alternative Splicing Variation in Human Populations. Am. J. Hum. Genet. 102, 11–26 (2018).

4. Wang, Z. & Burge, C. B. Splicing regulation: from a parts list of regulatory elements to an integrated splicing code. RNA N. Y. N 14, 802–813 (2008).

5. Scotti, M. M. & Swanson, M. S. RNA mis-splicing in disease. Nat. Rev. Genet. 17, 19–32 (2016).

6. Xiong, H. Y. et al. The human splicing code reveals new insights into the genetic determinants of disease. Science 347, 1254806 (2015).

7. Rosenberg, A. B., Patwardhan, R. P., Shendure, J. & Seelig, G. Learning the sequence determinants of alternative splicing from millions of random sequences. Cell 163, 698–711 (2015).

8. Cheng, J. et al. MMSplice: modular modeling improves the predictions of genetic variant effects on splicing. Genome Biol. 20, 48 (2019).

9. Jaganathan, K. et al. Predicting Splicing from Primary Sequence with Deep Learning. Cell 176, 535–548.e24 (2019).

10. MacArthur, D. G.et al. Guidelines for investigating causality of sequence variants in human disease. Nature 508, 469–476 (2014).

11. Richards, S. et al. Standards and Guidelines for the Interpretation of Sequence Variants: A Joint Consensus Recommendation of the American College of Medical Genetics and Genomics and the Association for Molecular Pathology. Genet. Med. Off. J. Am. Coll. Med. Genet. 17, 405–424 (2015).

12. Jian, X., Boerwinkle, E. & Liu, X. In silico prediction of splice-altering single nucleotide variants in the human genome. Nucleic Acids Res. 42, 13534–13544 (2014).

13. Sun, Y. et al. Next-generation diagnostics: gene panel, exome, or whole genome? Hum. Mutat. 36, 648–655 (2015).

14. Cummings, B. B.et al. Improving genetic diagnosis in Mendelian disease with transcriptome sequencing. Sci. Transl. Med. 9, eaal5209 (2017).

15. Kremer, L. S.et al. Genetic diagnosis of Mendelian disorders via RNA sequencing. Nat. Commun. 8, 15824 (2017).

16. Frésard, L. et al. Identification of rare-disease genes using blood transcriptome sequencing and large control cohorts. Nat. Med. 25, 911–919 (2019).

17. Gonorazky, H. D.et al. Expanding the Boundaries of RNA Sequencing as a Diagnostic Tool for Rare Mendelian Disease. Am. J. Hum. Genet. 104, 466–483 (2019).

18. Li, Y. I.et al. Annotation-free quantification of RNA splicing using LeafCutter. Nat. Genet. 50, 151–158 (2018).

19. Vincent, P., Larochelle, H., Bengio, Y. & Manzagol, P.-A. Extracting and Composing Robust Features with Denoising Autoencoders. in Proceedings of the 25th International Conference on Machine Learning 1096–1103 (ACM, 2008). doi:10.1145/1390156.1390294.

20. GTEx Consortium. The Genotype-Tissue Expression (GTEx) pilot analysis: multitissue gene regulation in humans. Science 348, 648–660 (2015).

21. Katz, Y., Wang, E. T., Airoldi, E. M. & Burge, C. B. Analysis and design of RNA sequencing experiments for identifying isoform regulation. Nat. Methods 7, 1009–1015 (2010).

22. Pervouchine, D. D., Knowles, D. G. & Guigó, R. Intron-centric estimation of alternative splicing from RNA-seq data. Bioinformatics 29, 273–274 (2013).

23. Li, X. et al. The impact of rare variation on gene expression across tissues. Nature 550, 239–243 (2017).

24. Ferraro, N. M.et al. Diverse transcriptomic signatures across human tissues identify functional rare genetic variation. bioRxiv 786053 (2019) doi:10.1101/786053.

25. Harrow, J. et al. GENCODE: The reference human genome annotation for The ENCODE Project. Genome Res. 22, 1760–1774 (2012).

26. Holm, S. A Simple Sequentially Rejective Multiple Test Procedure. Scand. J. Stat. 6, 65–70 (1979).

27. Benjamini, Y. & Yekutieli, D. The control of the false discovery rate in multiple testing under dependency. Ann. Stat. 29, 1165–1188 (2001).

28. Brechtmann, F. et al. OUTRIDER: A Statistical Method for Detecting Aberrantly Expressed Genes in RNA Sequencing Data. Am. J. Hum. Genet. 103, 907–917 (2018).

29. Karczewski, K. J.et al. Variation across 141,456 human exomes and genomes reveals the spectrum of loss-of-function intolerance across human protein-coding genes. bioRxiv 531210 (2019) doi:10.1101/531210.

30. Eilbeck, K. et al. The Sequence Ontology: a tool for the unification of genome annotations. Genome Biol. 6, R44 (2005).

31. McLaren, W. et al. The Ensembl Variant Effect Predictor. Genome Biol. 17, 122 (2016).

32. Drexler, H. L., Choquet, K. & Churchman, L. S. Human co-transcriptional splicing kinetics and coordination revealed by direct nascent RNA sequencing. bioRxiv 611020 (2019) doi:10.1101/611020.

33. Houtkooper, R. H.et al. The enigmatic role of tafazzin in cardiolipin metabolism. Biochim. Biophys. Acta 1788, 2003–2014 (2009).

34. Landrum, M. J.et al. ClinVar: improving access to variant interpretations and supporting evidence. Nucleic Acids Res. 46, D1062–D1067 (2018).

35. Ferri, L. et al. When silence is noise: infantile-onset Barth syndrome caused by a synonymous substitution affecting TAZ gene transcription. Clin. Genet. 90, 461–465 (2016).

36. Huber, W. et al. Orchestrating high-throughput genomic analysis with Bioconductor. Nat. Methods 12, 115–121 (2015).

37. R Core Team. R: A Language and Environment for Statistical Computing. (R Foundation for Statistical Computing, 2019).

38. Aicher, J. K., Jewell, P., Vaquero-Garcia, J., Barash, Y. & Bhoj, E. J. Mapping RNA splicing variations in clinically-accessible and non-accessible tissues to facilitate Mendelian disease diagnosis using RNA-seq. bioRxiv 727586 (2019) doi:10.1101/727586.

39. Casper, J. et al. The UCSC Genome Browser database: 2018 update. Nucleic Acids Res. 46, D762–D769 (2018).

40. Dobin, A. et al. STAR: ultrafast universal RNA-seq aligner. Bioinforma. Oxf. Engl. 29, 15–21 (2013).

41. Lawrence, M. et al. Software for computing and annotating genomic ranges. PLoS Comput. Biol. 9, e1003118 (2013).

42. Liao, Y., Smyth, G. K. & Shi, W. The R package Rsubread is easier, faster, cheaper and better for alignment and quantification of RNA sequencing reads. Nucleic Acids Res. 47, e47 (2019).

43. Braunschweig, U. et al. Widespread intron retention in mammals functionally tunes transcriptomes. Genome Res. 24, 1774–1786 (2014).

44. Khodor, Y. L. et al. Nascent-seq indicates widespread cotranscriptional pre-mRNA splicing in Drosophila. Genes Dev. 25, 2502–2512 (2011).

45. Tilgner, H. et al. Deep sequencing of subcellular RNA fractions shows splicing to be predominantly co-transcriptional in the human genome but inefficient for lncRNAs. Genome Res. 22, 1616–1625 (2012).

46. Stacklies, W., Redestig, H., Scholz, M., Walther, D. & Selbig, J. pcaMethods—a bioconductor package providing PCA methods for incomplete data. Bioinformatics 23, 1164–1167 (2007).

47. Byrd, R., Lu, P., Nocedal, J. & Zhu, C. A Limited Memory Algorithm for Bound Constrained Optimization. SIAM J. Sci. Comput. 16, 1190–1208 (1995).

48. Zhou, X., Lindsay, H. & Robinson, M. D. Robustly detecting differential expression in RNA sequencing data using observation weights. Nucleic Acids Res. 42, e91 (2014).

49. Yee, T. W. Vector Generalized Linear and Additive Models: With an Implementation in R. (Springer New York, 2015). doi:10.1007/978-1-4939-2818-7_1.

