## Supplementary Materials for "Detection of aberrant splicing events in RNA-seq data with FRASER"

### 1 Supplemental Figures

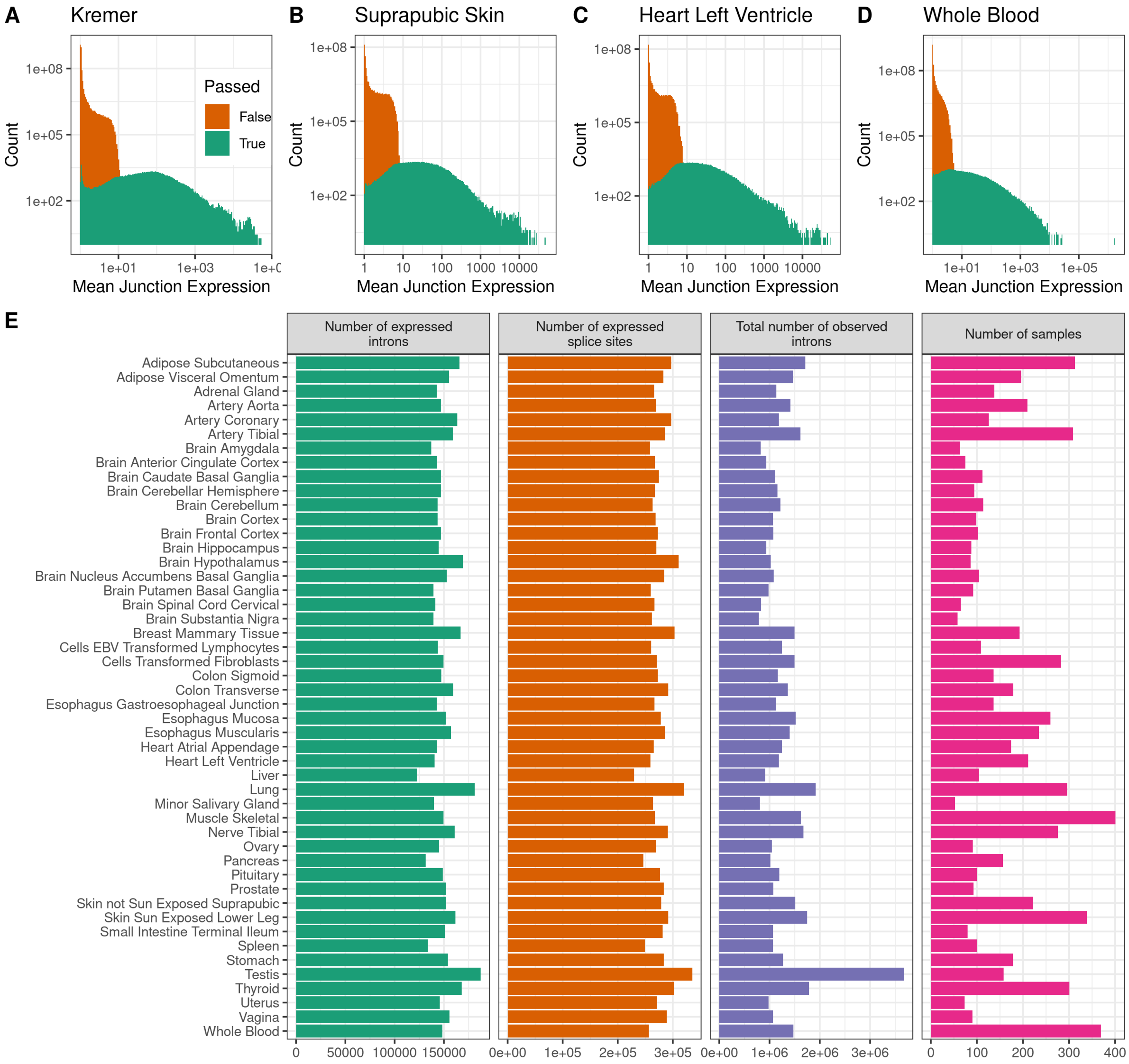


**Figure S1**: **Filtering of introns.** **(A)** Histogram of the raw intron coverage per sample-intron pair for the Kremer data set grouped according to the intron filter status. Green indicates that the intron passed the filter and orange indicates that the intron was filtered out. **(B-D)** Same as A, but for different tissues in the GTEx data set. **(E)** Barplots of the number of introns passed the filtering, splice sites passed the filtering, observed introns, and samples per tissue within the GTEx data set used in this study.

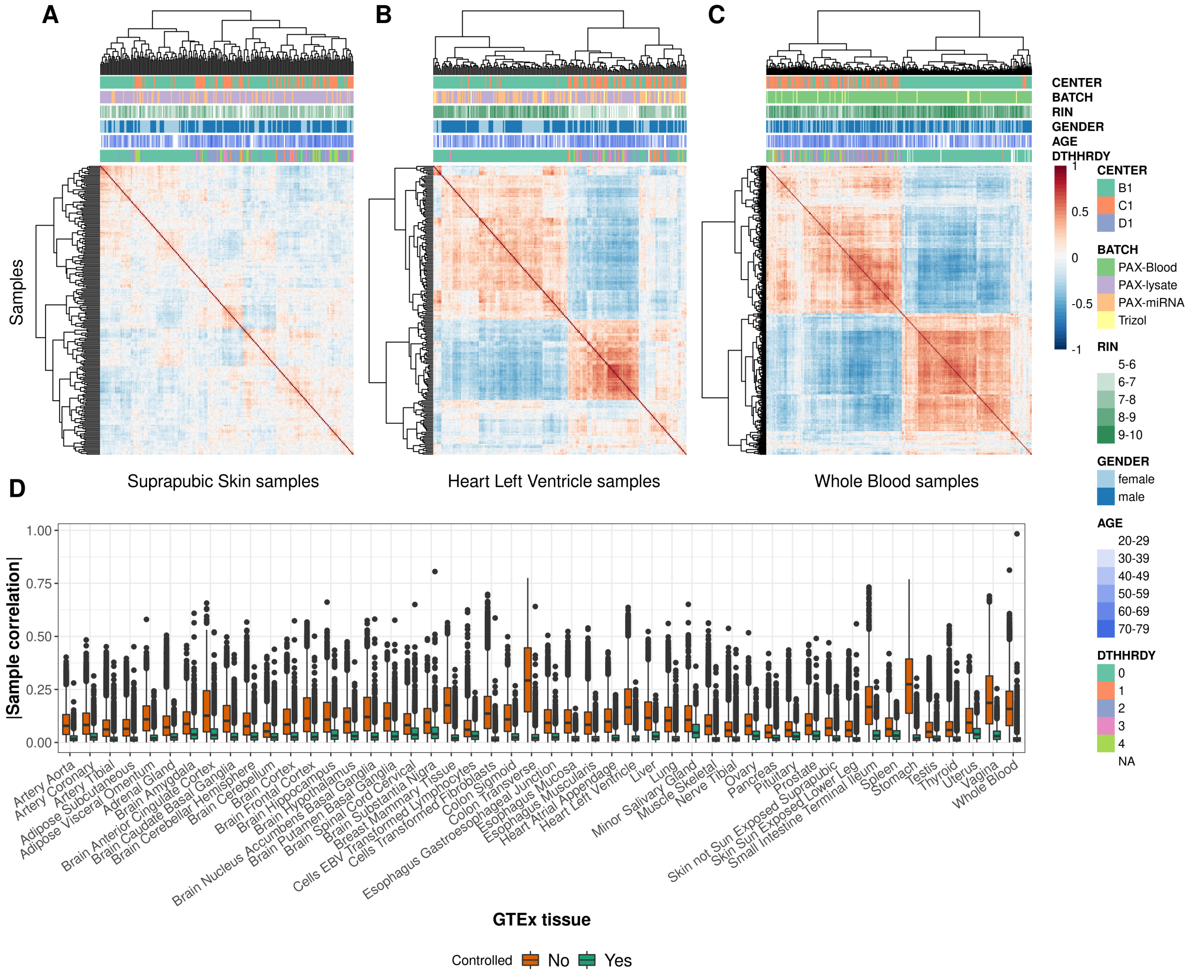


**Figure S2**: **Tissue-specific correlation structure for $\psi_3$. (A-C)** Intron-centered and logit-transformed $\psi_3$ of the 10,000 most variable introns clustered by samples (columns and rows) for three representative GTEx tissues: suprapubic skin (A), left ventricle heart (B), and whole blood (C). Red and blue depict high and low intron usage, respectively. Colored horizontal tracks display sequencing center, batch, RNA integrity number (RIN), gender, age, and cause of death (DTHHRDY, Hardy scale classification) of the samples. **(D)** Boxplot of a bsolute values of between-sample correlations of row-centered logit-transformed $\psi_3$ for 48 GTEx tissues before (orange) and after (green) correction for the latent space.

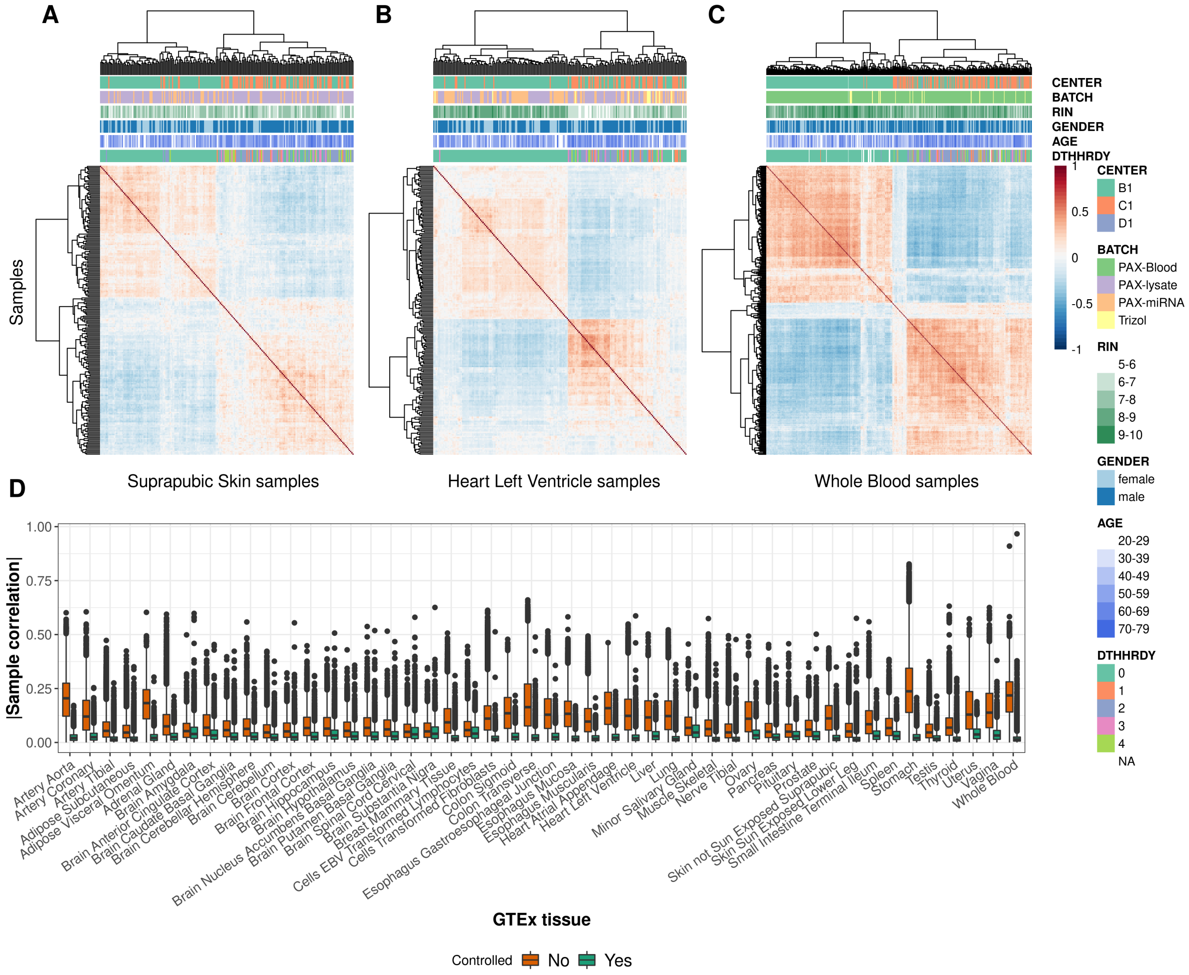


**Figure S3**: **Tissue-specific correlation structure for $\theta$.** **(A-C)** Splice-site-centered and logit-transformed $\theta$ of the 10,000 most variable splice sites clustered by samples (columns and rows) for three representative GTEx tissues: suprapubic skin (A), left ventricle heart (B), and whole blood (C). Red and blue depict high and low intron usage, respectively. Colored horizontal tracks display sequencing center, batch, RNA integrity number (RIN), gender, age, and cause of death (DTHHRDY, Hardy scale classification) of the samples. **(D)** Boxplot of a bsolute values of between-sample correlations of row-centered logit-transformed $\theta$ for 48 GTEx tissues before (orange) and after (green) correction for the latent space.

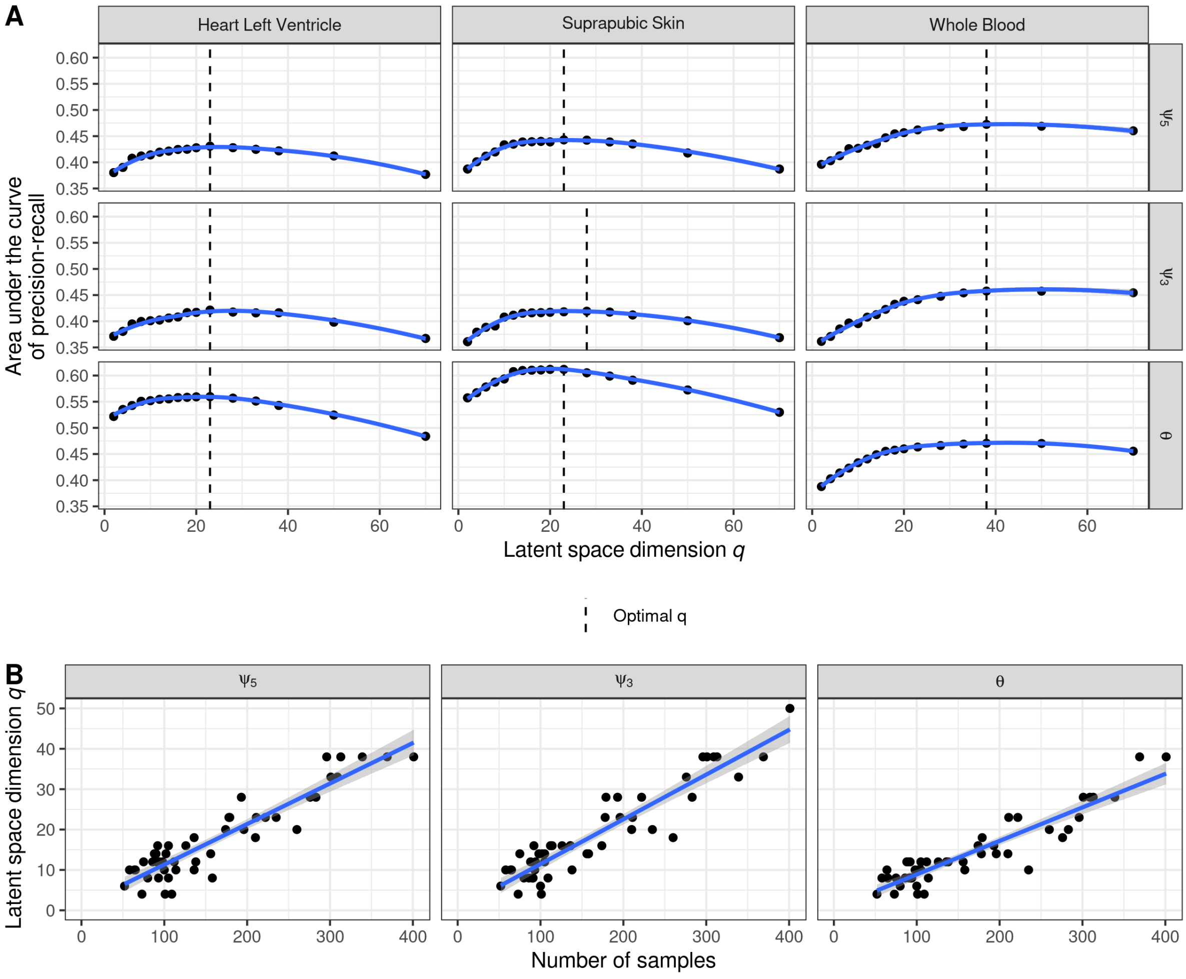


**Figure S4**: **Finding the optimal latent space dimension** $q$**. (A)** The area under the precision recall curve is plotted for a given latent space dimension $q$. The dashed line is the optimal $q$ for the given data set. The blue line is a local polynomial regression fit. The data is facet by the splicing metric (rows) and the three example datasets from GTEx (columns). **(B)** For each of the 48 GTEx tissues, the number of samples are plotted against the estimated latent space dimension. The data is stratified by the splicing metrics (columns). The blue line represents a linar regression fit and the gray band around it defines the 95% confidence interval of the fit.

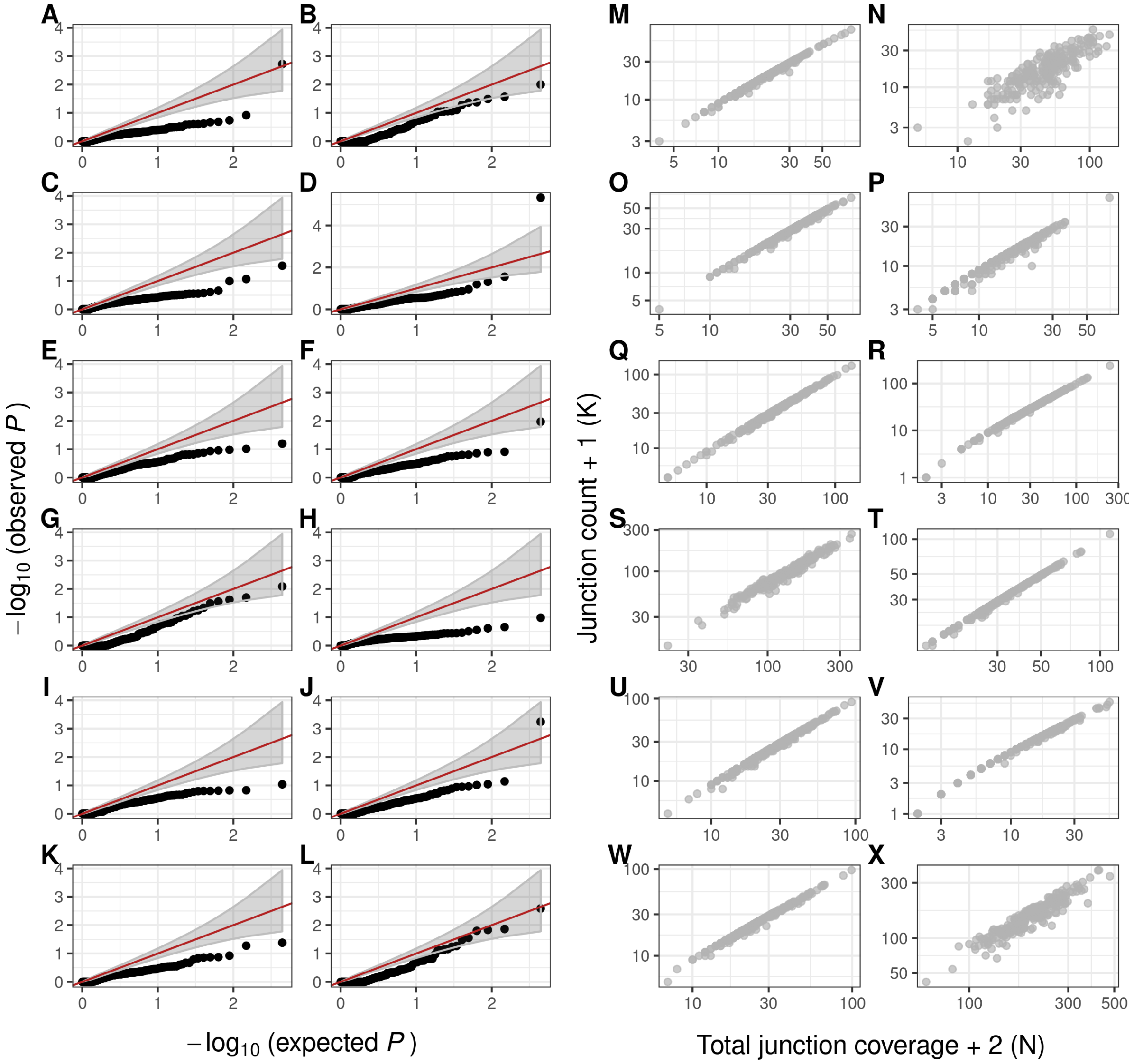


**Figure S5**: **Quantile-quantile plots and the corresponding count ratios. (A-L)** Quantile-quantile plots of randomly chosen introns based on the $\psi_5$ metric. **(M-X)** Expression plots of the number of split reads (K) over the total coverage (N) of the given donor site. The q-q plots correspond to the respective expression plot. The data is based on the suprapubic skin GTEx tissue.

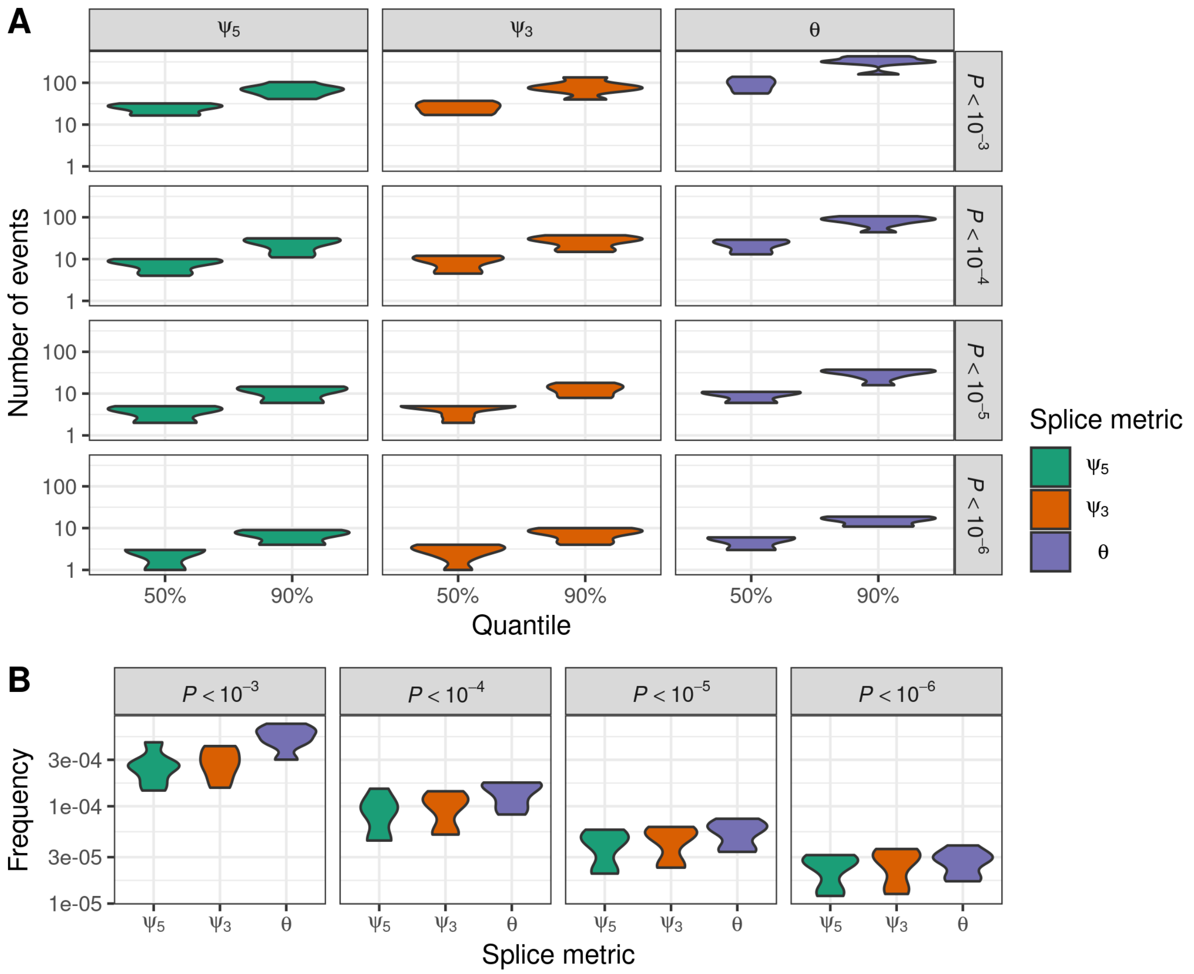


**Figure S6**: **Distribution of extreme beta-binomial *P* values across GTEx tissues. (A)** The distribution of the number of events (y-axis) having a more significant $P$ value than a given cutoff (rows) is plotted against the quantiles across samples within a tissue (x-axis, median and 90%). The data is stratified by the different splicing metrics (columns, with green, orange, and violette for $\psi_5$, $\psi_3$, and $\theta$, respectively). Each distribution is based on the 48 GTEx tissues. **(B)** Same as in A but the frequency (y-axis) of $P$ values being smaller than a given cutoff (columns) across a tissue is plotted per splicing metrics (x-axis).

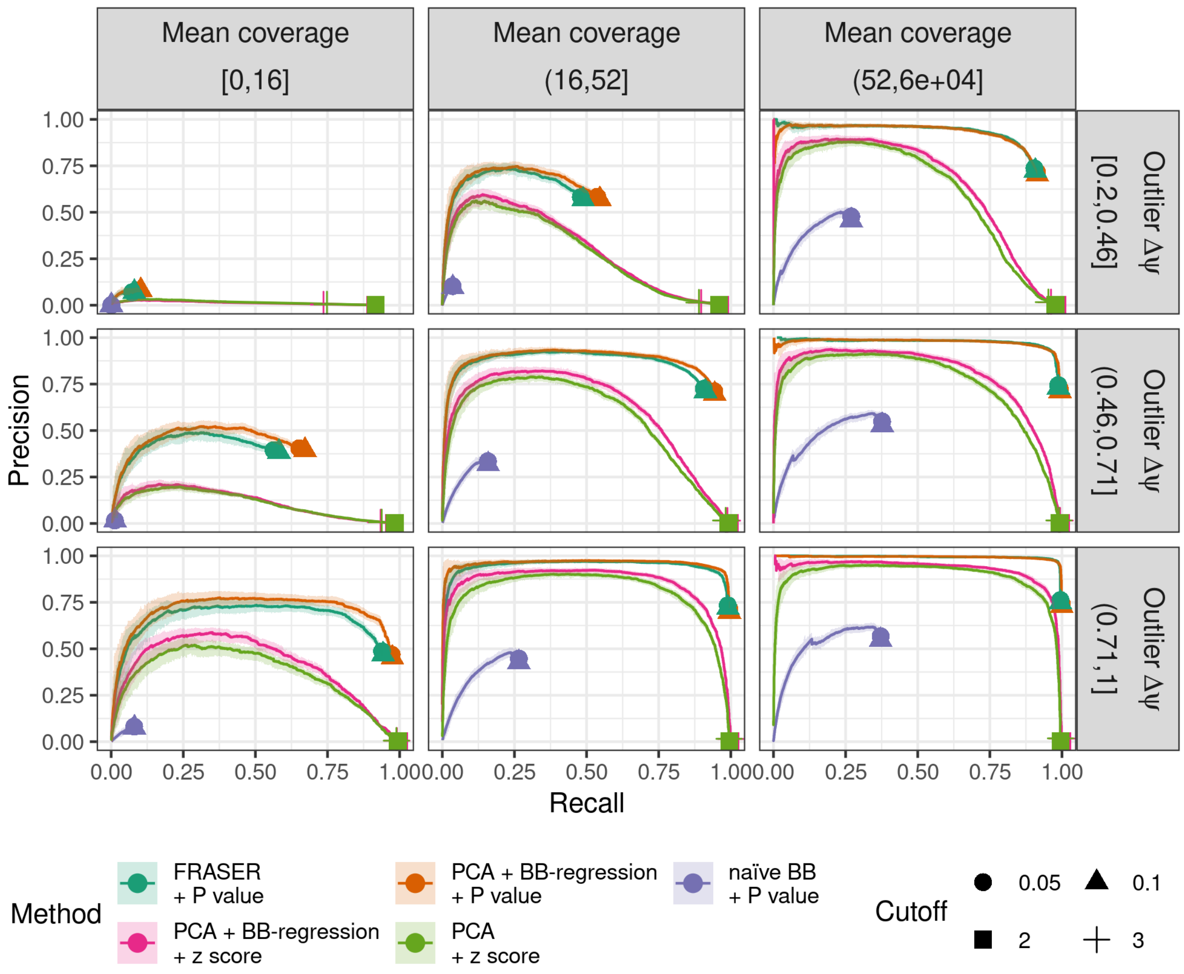


**Figure S7**: **Splicing outlier detection benchmark in GTEx for $\psi_3$.** The proportion of simulated outliers among reported outliers (precision, y-axis) plotted against the proportion of reported simulated outliers among all simulated outliers (recall, x-axis) for increasing beta-binomial $P$ values computed using count ratio expectations based on FRASER (green), a beta-binomial regression on the latent space (orange), or on raw count ratios (violette, naïve BB) and for decreasing absolute z scores ontop of a beta-binomial regression (pink) or PCA. Additionally, all events with $|\Delta\psi| < 0.1$ are ranked last. Plots are stratified equally by injected amplitudes ($\Delta\psi$, by row) and junction coverage (by column). The points indicate commonly applied cutoffs (FDR $< 0.1$ and $< 0.05$ and absolute z scores $> 2$ and $> 3$). Light ribbons around the curves depict 95% confidence bands estimated by bootstrapping.

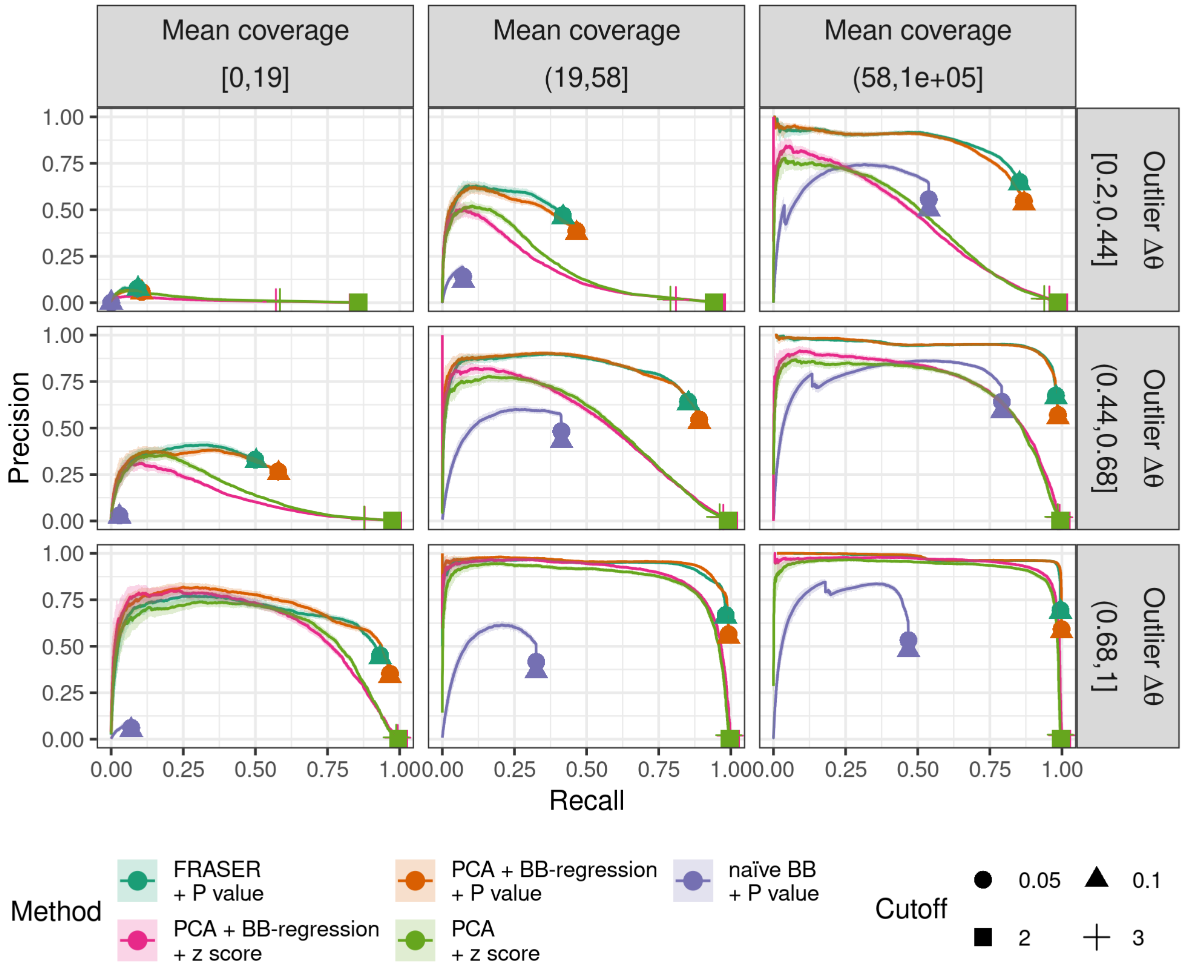


**Figure S8**: **Splicing outlier detection benchmark in GTEx for** $\theta$**.** Same as Figure **S7**, but based on the splicing efficiency metric $\theta$.

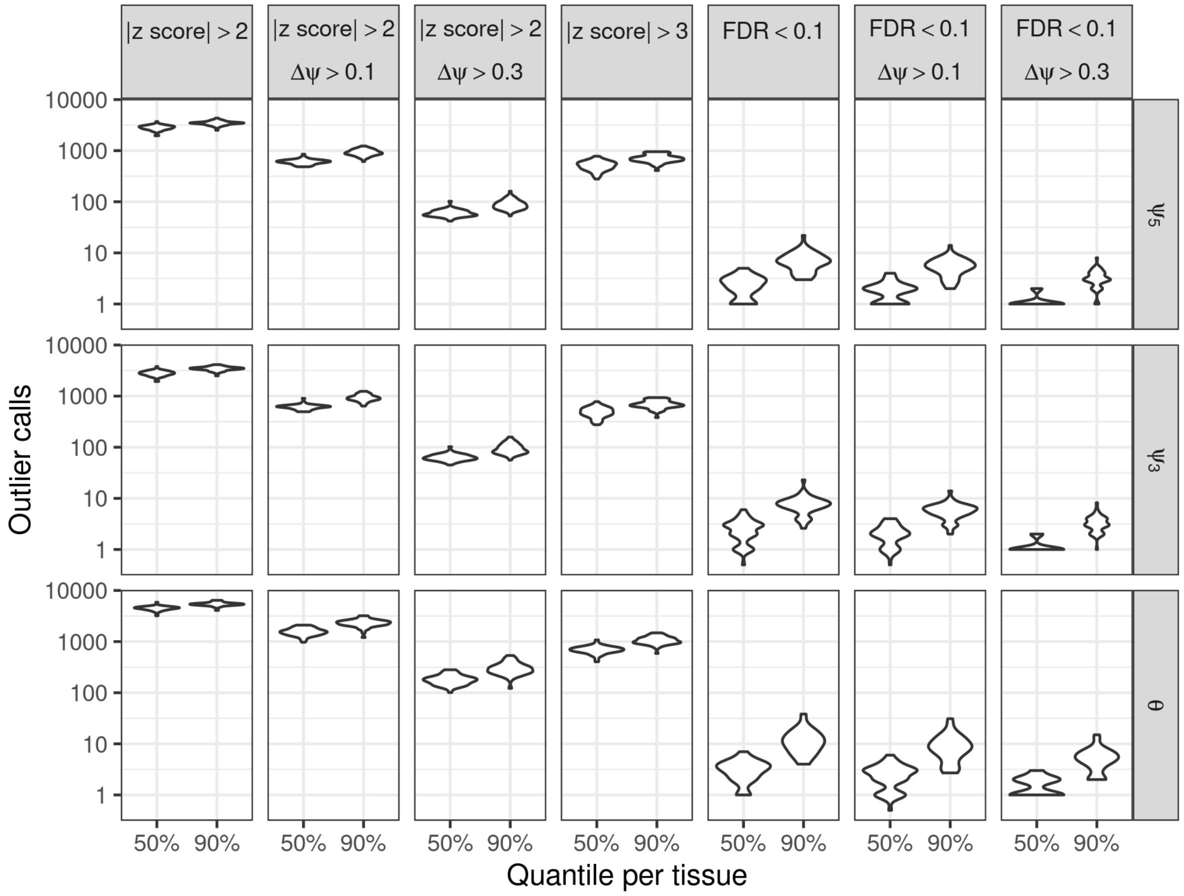


**Figure S9**: **Distribution of aberrant splicing events in GTEx.** Distribution of aberrant splicing events (y-axis) called using different cutoffs based on the FRASER normalization are plotted against two quantiles (rows, 50% and 90%) across samples in a tissue for all 48 GTEx tissues. The plot is stratified by the splicing metrics (rows) and commonly applied cutoffs (columns). Overall z score based cutoffs report in magnitudes more outlier than significant based and FDR controlled cutoff approaches.

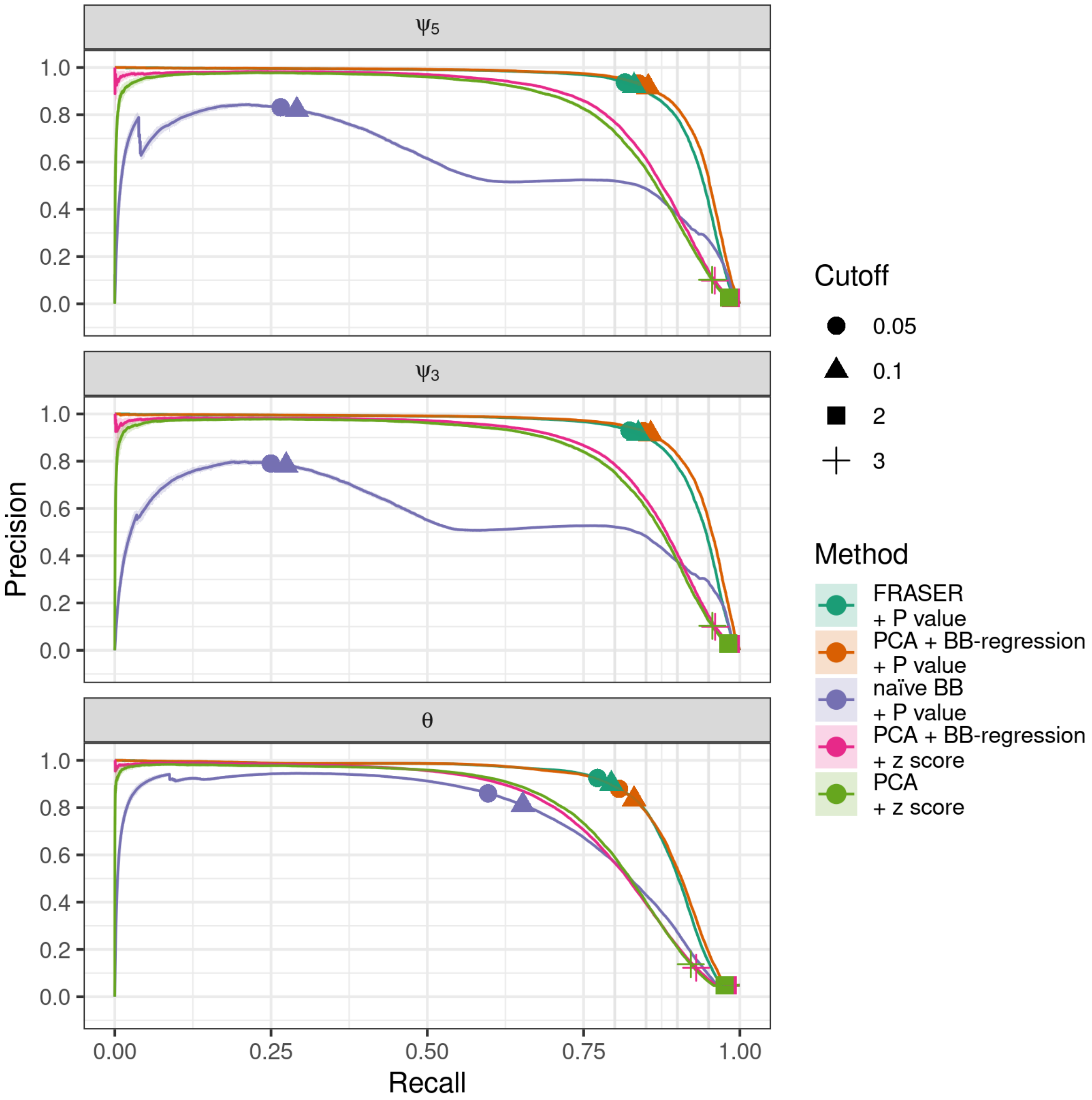


**Figure S10**: **Performance benchmark using artificially injected outliers.** The proportion of simulated outliers among reported outliers (precision, y-axis) plotted against the proportion of reported simulated outliers among all simulated outliers (recall, x-axis) for increasing beta-binomial $P$ values computed using count ratio expectations based on FRASER (green), a beta-binomial regression on the latent space (orange), or on raw count ratios (violette, naïve BB) and for decreasing absolute z scores ontop of a beta-binomial regression (pink) or PCA. The data is stratified by the different splice metrics: $\psi_5$, $\psi_3$, and $\theta$ (rows). The points indicate commonly applied cutoffs (FDR $< 0.1$ and $< 0.05$ and absolute z scores $> 2$ and $> 3$). Light ribbons around the curves depict 95% confidence bands estimated by bootstrapping.

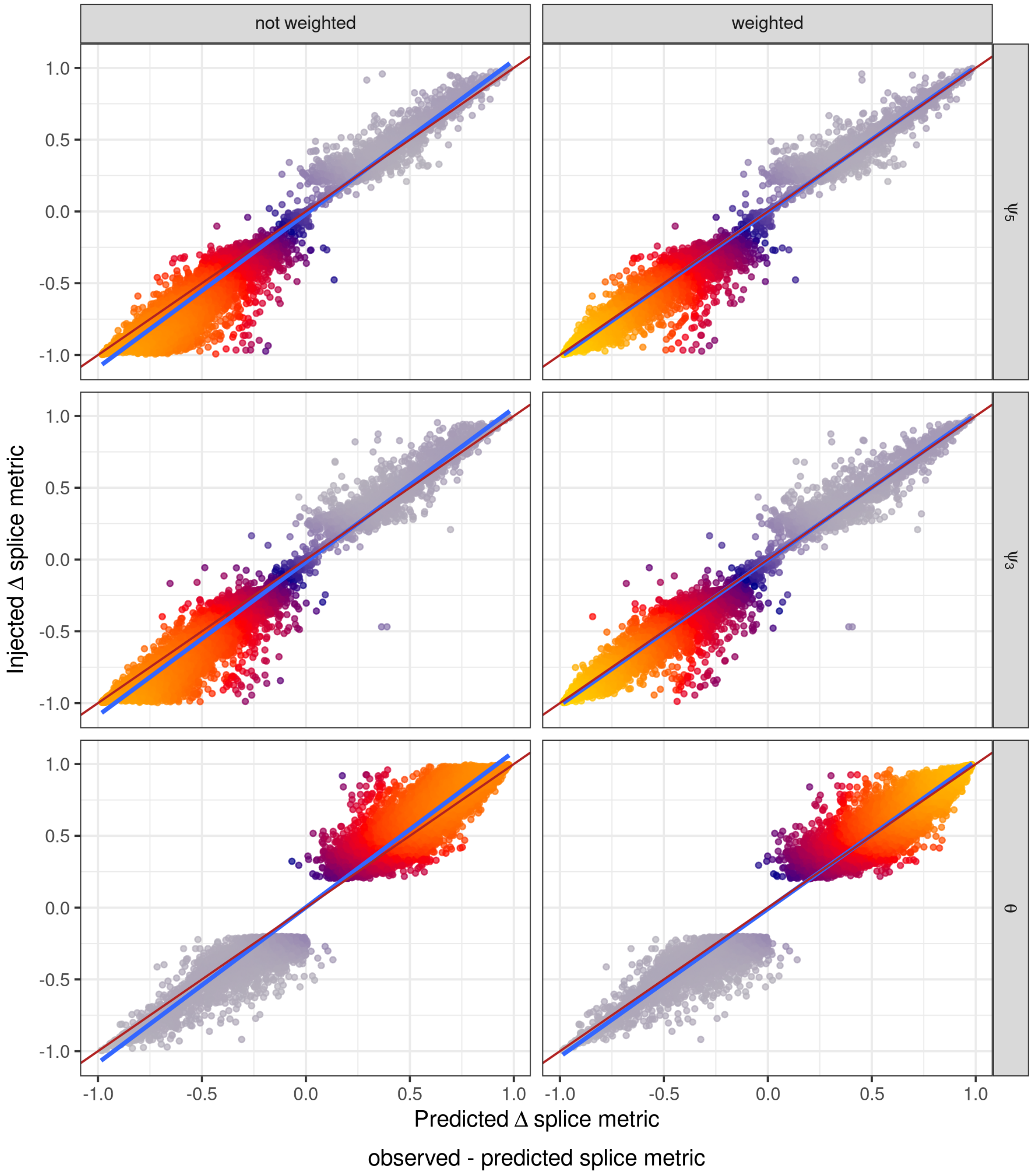


**Figure S11**: **Using a weighted beta-binomial loss for robustness against outlier data points.** For each injected outlier data point, the effect size of the injection (y-axis, $\Delta\bullet$) is plotted against the predicted difference (x-axis, observerd $\Delta\bullet -$ predicted $\Delta\bullet$) based on the beta-binomial regression fit. The plot is stratified by the three splicing metrics $\psi_5$, $\psi_3$, and $\theta$ (rows) and by loss function used in the beta-binomial regression (columns, non weighted and weighted). The blue line corresponds to a linear regression. The red line indicates the diagonal, which would be the perfect fit. The data is based on the suprapubic skin GTEx tissue.

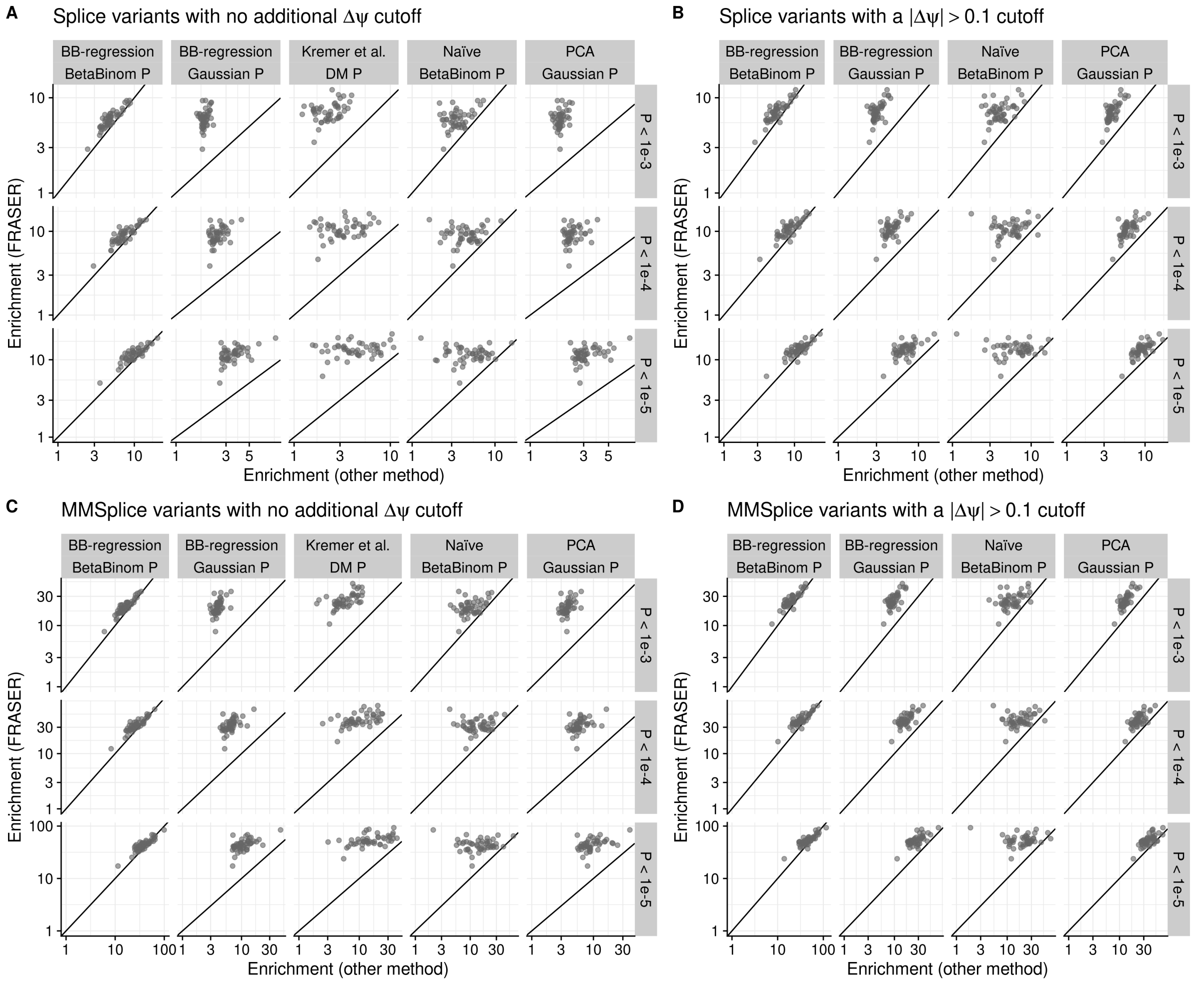


**Figure S12**: **Gene-based rare variant enrichment analysis.** **(A)** Enrichment using FRASER (y-axis) against enrichment using different aberrant splicing detection methods (x-axis, columns) for rare variants located in a splice region. The method applied are a beta-binomial regression on the latent space with (i) beta-binomial $P$ values and (ii) Gaussian $P$ values, (iii) Dirichlet-Multinomial $P$ values based on Kremer et al., (iv) nave beta-binomial $P$ values, and (v) PCA-based Gaussian $P$ values. The enrichment is calculated for different nominal $P$ value cutoffs (rows). Each dot represents a GTEx tissue. **(B)** The same as A but an additional $|\Delta\psi| > 0.1$ cutoff is applied on the aberrant splicing calles. **(C)** The same as A but based on rare variants predicted to afffect splicing by MMSplice. **(D)** The same as C but an additional $|\Delta\psi| > 0.1$ cutoff is applied on the aberrant splicing calles.

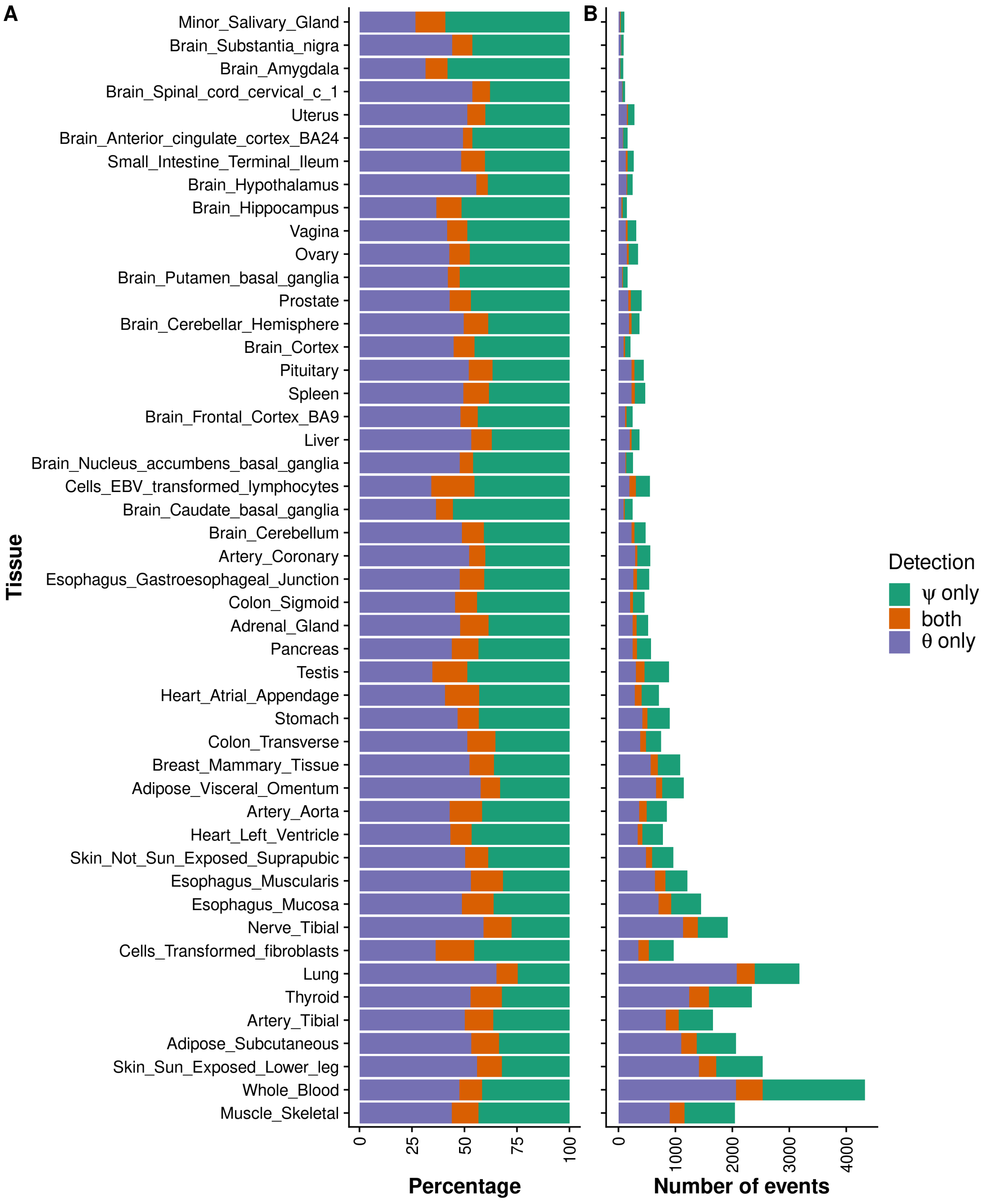


**Figure S13**: **Contribution of intron retention in aberrant splicing. (A)** Barplot of the percentage of aberrant splicing events on the gene level grouped by the detection metrics: alternative splicing only (violette, $\psi$ metric), intron retention (green, $\theta$ metric), and both (red). (B) Same as A but with the absolute number of detected events.

### 2 Supplemental Methods

#### 2.1 Fitting of the parameters

All notations are introduced in the Materials and Methods section.

##### Beta-binomial model

We use the following parameterization of the beta-binomial distribution:

$$P(k|n,\alpha,\beta) = \frac{\Gamma(n+1)}{\Gamma(k+1)\Gamma(n-k+1)} \frac{\Gamma(\alpha+k)\Gamma(\beta+n-k)}{\Gamma(\alpha+\beta+n)} \frac{\Gamma(\alpha+\beta)}{\Gamma(\alpha)\Gamma(\beta)},$$

where

$$\alpha = \mu\left(\frac{1-\rho}{\rho}\right) \text{ and } \beta = (\mu-1)\left(\frac{\rho-1}{\rho}\right).$$

The variance of the distribution is given by:

$$Var = \frac{\mu(1-\mu)(1+(n-1)\rho)}{n}.$$

##### The negative log-likelihood of the beta-binomial distribution

The negative log-likelihood (nll) of the model is given by:

$$\begin{aligned}
\text{nll} = & -\sum_{ij}\log(\Gamma(n_{ij}+1)) + \sum_{ij} log(\Gamma(k_{ij}+1)) + \sum_{ij}\log(\Gamma(n_{ij}-k_{ij}+1)) \\
& -\sum_{ij}\log(\Gamma(\alpha_{ij}+k_{ij})) - \sum_{ij}\log(\Gamma(\beta_{ij}+n_{ij}-k_{ij})) + \sum_{ij}\log(\Gamma(\alpha_{ij}+\beta_{ij}+n_{ij})) \\
& +\sum_{ij}\log(\Gamma(\alpha_{ij})) + \sum_{ij}\log(\Gamma(\beta_{ij})) - \sum_{ij}\log(\Gamma(\alpha_{ij}+\beta_{ij})).
\end{aligned}$$

For the optimization of the model only the terms of nll that are dependent on $\mathbf{W}_e$ and $\mathbf{W}_d$ need to be considered. Since

$$\alpha + \beta = \frac{1-\rho}{\rho}$$

is independent of $\mu$ and therefore independent of $\mathbf{W}_e$ and $\mathbf{W}_d$, we do not have to consider the terms containing $\alpha+\beta$ and yield the following truncated form of the negative log likelihood, with pseudocounts of 1 and 2 added to $k$ and $n$, respectively:

$$\text{nll}_{\mathbf{W}} = \sum_{ij}\log(\Gamma(\alpha_{ij})) + \sum_{ij}\log(\Gamma(\beta_{ij})) \tag{1}$$

$$-\sum_{ij}\log(\Gamma(\alpha_{ij}+k_{ij}+1)) - \sum_{ij}\log(\Gamma(\beta_{ij}+n_{ij}-k_{ij}+1)). \tag{2}$$

In the following $y_{ij}$ is an element of $\mathbf{Y}$ defined as:

$$\mathbf{Y} = \widetilde{\mathbf{X}}\mathbf{W}_e\mathbf{W}_d + \mathbf{b}, \tag{3}$$

where the element $\widetilde{x}_{ij}$ of the matrix $\widetilde{\mathbf{X}}$ is given by:

$$\begin{aligned}
\widetilde{x}_{ij} &= x_{ij} - \bar{x}_j, \\
x_{ij} &= \operatorname{logit}\left(\frac{k_{ij}+1}{n_{ij}+2}\right), \operatorname{logit}(a) &= \log\frac{a}{1-a}.
\end{aligned}$$

The expectations $\mu_{ij}$ are then modeled by:

$$\mu_{ij} = \sigma\left(y_{ij}\right) = \frac{e^{y_{ij}}}{1+e^{y_{ij}}}$$

Hence, $\text{nll}_{\mathbf{W}}$ can be rewritten as:

$$\begin{aligned}
\text{nll}_{\mathbf{W}} = &\sum_{ij} \log\left(\Gamma\left(\frac{e^{y_{ij}}}{1+e^{y_{ij}}}\frac{1-\rho_{ij}}{\rho_{ij}}\right)\right) \\
&+ \sum_{ij} \log\left(\Gamma\left(\left(\frac{e^{y_{ij}}}{1+e^{y_{ij}}}-1\right)\frac{\rho_{ij}-1}{\rho_{ij}}\right)\right) \\
&- \sum_{ij} \log\left(\Gamma\left(\frac{e^{y_{ij}}}{1+e^{y_{ij}}}\frac{1-\rho_{ij}}{\rho_{ij}} + k_{ij} + 1\right)\right) \\
&- \sum_{ij} \log\left(\Gamma\left(\left(\frac{e^{y_{ij}}}{1+e^{y_{ij}}}-1\right)\frac{\rho_{ij}-1}{\rho_{ij}} + n_{ij} - k_{ij} + 1\right)\right)
\end{aligned}$$

We use L-BFGS[1] as implemented in *optim* to fit the autoencoder model as described in Methods.

##### Update of the encoder and decoder matrix

The updating of the matrix $\mathbf{W}_d$ is performed intron-wise where as the encoder matrix $\mathbf{W}_e$ is performed on the full matrix. For each update step, the intron-wise or matrix-wise average negative log likelihood is minimized. To not run into convergence issues or numerical instability of the digamma function, we estimate the value of the digamma function $\psi$ if not $-35 < y_{ij} < 30$. From Equation 1 and Equation 3, we obtain the gradients:

$$\begin{aligned}
\frac{d\text{nll}}{d\mathbf{W}_e} &= \widetilde{\mathbf{X}}^T\mathbf{A}\mathbf{W}_d + \widetilde{\mathbf{X}}^T\mathbf{B}\mathbf{W}_d - \widetilde{\mathbf{X}}^T\mathbf{C}\mathbf{W}_d - \widetilde{\mathbf{X}}^T\mathbf{D}\mathbf{W}_d \\
\frac{d\text{nll}}{d\mathbf{W}_d} &= \mathbf{A}^T\widetilde{\mathbf{X}}\mathbf{W}_e + \mathbf{B}^T\widetilde{\mathbf{X}}\mathbf{W}_e - \mathbf{C}^T\widetilde{\mathbf{X}}\mathbf{W}_e - \mathbf{D}^T\widetilde{\mathbf{X}}\mathbf{W}_e \\
\frac{d\text{nll}}{db_j} &= \sum_i a_{ij} + b_{ij} - c_{ij} - d_{ij}
\end{aligned}$$

,

where the components of the matrices $\mathbf{A}, \mathbf{B}, \mathbf{C}$ and $\mathbf{D}$ are computed by:

$$
\begin{aligned}
a_{ij} &= \psi\left(\frac{e^{y_{ij}}}{1+e^{y_{ij}}} \cdot r_{ij}\right) \cdot r_{ij} \cdot v_{ij} \\
b_{ij} &= \psi\left(\left(\frac{e^{y_{ij}}}{1+e^{y_{ij}}} - 1\right) \cdot (-r_{ij}\right) \cdot (-r_{ij}) \cdot v_{ij} \\
c_{ij} &= \psi\left(\frac{e^{y_{ij}}}{1+e^{y_{ij}}} \cdot r_{ij} + k_{ij} + 1\right) \cdot r_{ij} \cdot v_{ij} \\
d_{ij} &= \psi\left(\left(\frac{e^{y_{ij}}}{1+e^{y_{ij}}} - 1\right) \cdot (-r_{ij} + n_{ij} - k_{ij} + 1\right) \cdot (-r_{ij}) \cdot v_{ij} \\
v_{ij} &= \frac{e^{y_{ij}}}{\left(1+e^{y_{ij}}\right)^2} \\
r_{ij} &= \frac{1-\rho_j}{\rho_j}
\end{aligned}
$$

.

### References


[1] Byrd, R., Lu, P., Nocedal, J., and Zhu, C. (1995). A Limited Memory Algorithm for Bound Constrained Optimization. SIAM Journal on Scientific Computing *16*, 1190–1208.
